# The information signature of diverging lineages

**DOI:** 10.1101/2021.08.30.458276

**Authors:** Douglas G. Moore, Matheo Morales, Austin R. Biddy, Sara I. Walker, Greer A. Dolby

## Abstract

Divergence and speciation are the generators of biodiversity on Earth. Yet, we still seek to understand why species are unevenly distributed across taxa and geographic settings. This is partly because we lack a way to directly compare aspects of population divergence across unrelated species and an integrative quantitative framework for how geographical setting relates to genetic divergence. Here we postulate that, based on communications theory, there is a communications channel between Earth’s surface and the structuring of genomic information among populations. Each species is a separate receiver of this message. Organism traits determine the clarity of that channel. Calculating the partial information decomposition (PID) of genetic metrics under different evolutionary scenarios can measure the signature of this communication. We show *in silico* how the informational decomposition of 97,200 genetic lattices varies as lineages diverge. The decomposed nodes of Tajima’s D, θ_W_, and π show strong context-dependent information signatures, while F_ST_ was least informative. Specific decompositions detected whether lineages divergence with or without gene flow better than current SFS-based tools. The study of information in an evolutionary genomic and landscape context suggests it may be a helpful framework and for understanding the diversity of life across Earth and their coevolution.

## INTRODUCTION

Population differentiation and speciation are core processes within evolutionary biology because they are the only ways in which biodiversity is generated *de novo.* Accordingly, a tremendous amount is known about the mechanisms and context of speciation in individual systems, such as *Heliconius* butterflies (Martin et al., 2013; Merrill et al., 2015; Rougemont et al., 2023; Supple et al., 2015), stickleback fishes (Hendry et al., 2009; Liu et al., 2022; Marques et al., 2017; Ravinet et al., 2018) and monkeyflowers (Chase et al., 2017; Stankowski et al., 2019, 2023). Polyploidization, sexual selection, and ecological adaptation are just some modes by which species have diverged (Heslop-Harrison et al., 2023; Marques et al., 2017). Yet, core questions remain about how often lineages diverge in complete isolation (vicariance) or with gene flow (parapatry, sympatry). While speciation via sexual selection or ecological adaptation may be expected to occur in the presence of gene flow, there remains intense interest in the role geological, climatic, and geographical settings play in mediating or promoting speciation (Antonelli, 2017; Antonelli et al., 2018; Dolby, 2021; Rahbek et al., 2019; Ribas et al., 2022) particularly in the nascent field of geogenomics (Dolby et al., 2022). These settings may be expected to impact gene flow in different ways. A flooded landscape may entirely reduce gene flow for some species, while different climatic patterns may lead to low levels of ongoing gene flow as populations adapt to different conditions.

Speciation genomic studies must often focus on few, closely related species, yet understanding the frequency at which different speciation contexts occur in nature requires integrating the results of individual studies to yield broader patterns. Yet it is well known that changes in effective population size, recombination rate (Samuk et al., 2017), and other species-specific effects directly influence the measures of population diversity and divergence (e.g., π, F_ST_) we use to characterize population differentiation and evolutionary history. These traits vary dramatically across species (ArayaCDonoso et al., 2022), making it difficult to compare characteristics of diverging populations across unrelated species in the same geographical setting (Figure 1). For example, river networks are long hypothesized to structure populations, leading to allopatric speciation (see Naka et al., 2022). Answering this question might require comparing levels of differentiation using pairwise F_ST_ among river-isolated populations in a broad array of taxa (e.g., birds, insects, herbs), but F_ST_ is affected by species-specific characteristics. Analyzing the site frequency spectrum (SFS; Xue & Hickerson, 2015) offers a good option, but has some limitations because different histories can produce similar geometric distributions (Myers et al., 2008). Therefore, to uncover more generalized patterns about what settings lead to speciation requires comparing speciation history amongst disparate taxa, and new analytical frameworks may be helpful to achieving this.

**Figure 1.**
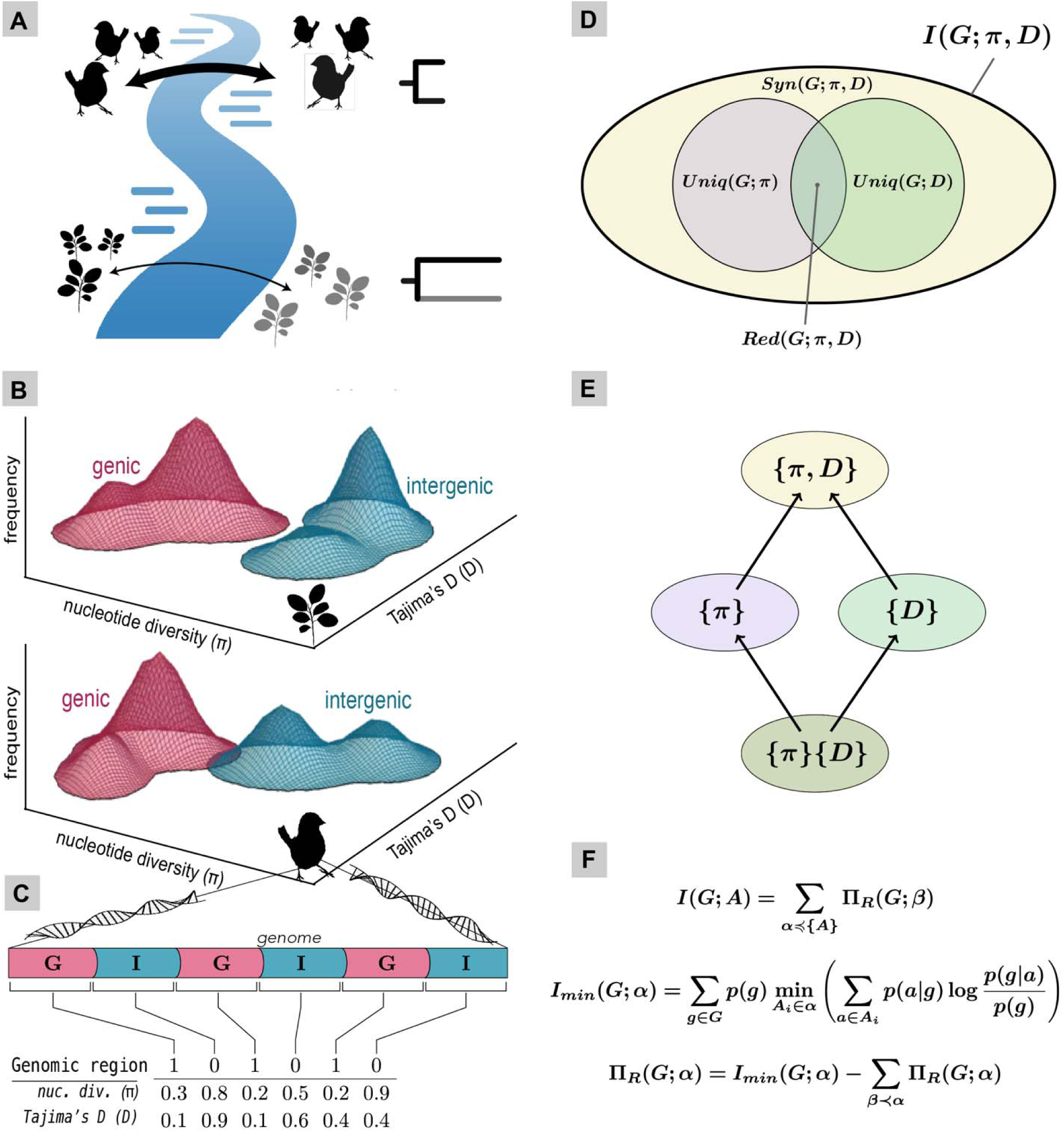
Conceptualization of partial information decomposition (PID) applied to genomic data (gPID). **(A)** Organisms may be affected differently by the same extrinsic events (e.g., formation of a river). **(B)** Genic and intergenic parts of the genome are differentially shaped by extrinsic (e.g., barrier to gene flow) and intrinsic (e.g., selection) forces that can be quantified with genetic metrics (e.g., F_ST_, Tajima’s D), the values of which will vary by taxon. **(C)** We calculated common metrics (π-nucleotide diversity, D-Tajima’s D) in genic and intergenic regions of the genome on simulated data and **(D, E)** calculated mutual information on these genetic statistics which is decomposed over a lattice into unique, redundant, synergistic information terms given by **(F)**.

DNA is well accepted as an informational molecule: it encodes “directions” for both RNA and protein synthesis, the redundancy of the duplex’s antiparallel strands forms the basis for DNA replication and recombination-dependent DNA repair synthesis, and that sequence is nearly perfectly transmitted to offspring save for the effects of meiotic recombination and new mutations. Yet DNA, in the form of genetic variation across geographical space, also records features of the landscape extrinsic to the organism. Simplistically, say at T_0_ we observe a landscape with unvarying climate, hydrography, and topography at the kilometer scale; individuals freely breed across the region. If we analyze the genomes of individuals at T_0_ there is no population structure save for perhaps isolation by distance (IBD). At T_1_ a drainage network or set of topography develops that limits individuals’ ability to move across that feature. At T_2_, many generations later, we sample the genomes of individuals across that landscape; we would discover genetic variation is now structured over space beyond the effects of IBD: the landscape has “communicated” some form of *information* which is now recorded in DNA of that species at the population level. Even if we never observe the landscape, we can know something about the feature that formed through the patterns of population genomic variation, perhaps its approximate age, location, or size, by observing DNA sequences alone. This (implicitly) was the theoretical basis for geogenomics (Baker et al., 2014) and is, arguably, part of the rationale behind the field of biogeography (Crisci, 2001).

In the framing of communications theory, this means there is an informational channel open between Earth’s landscape on thousand-to-million-year timescales (i.e., “mesoscale” *sensu* Dolby et al., 2022) and the population genomic patterns recorded in the structuring of genomic variants over geographical space. Every species or evolutionary lineage is an alternate receiver of this single communication. The characteristics of the organism (e.g., dispersal mode, generation time) mediate the clarity of its channel. That is, whether populations evolve responsively to the landscape: a “noiseless” channel; or, whether populations are unaffected: a “noisy” channel. In this study we document a proof-of-concept of that channel through simulation of large genome sequences and quantification of information communicated to the DNA of those populations using common genetic metrics.

We will give a brief background to understand what information is because the concept can be elusive and the term itself misleading. In the context of information theory, information is distilled down into a variety of measures of how much the uncertainty of one or more variables of interest is reduced by observations of other variables. For example, perhaps we want to learn (gain information) about whether it recently rained by observing whether the grass is wet. If the grass is dry, you can be reasonably certain that it did not rain (though it could have rained and then dried). However, the observation that the grass is wet reduces your uncertainty by less than the observation that the grass is dry because other factors can explain damp grass: dew, the running of a sprinkler system, etc. By observing whether the grass is wet (the source variable), one can gain information about whether or not it rained (the target variable). Yet, it only provides partial information. Additional observations, say overnight temperatures, the dew point, and whether or not your sprinkler system is broken can further reduce your uncertainty. Other observations can change the value (or weight) of those observations, for example knowing if it is the rainy season or not. Meanwhile other observations, such as whether your newspaper was delivered or not, offer little to no information about whether it rained. Information theory, at its core, provides a framework for quantifying these concepts.

The fundamental quantity of information, entropy, was introduced by Claude Shannon and provides a measure of the amount of uncertainty in a random variable (Shannon, 1948). It is the basis of the Shannon Index, H, which measures the entropy of an ecosystem based on species richnesses. In the context of standard statistics, it is often closely related to variance, e.g., the entropy of a normal distribution is 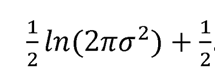. Reducing uncertainty in a random variable is similar to reducing the variance of the variable’s underlying distribution. In that same work, Shannon went on to present a formal conception of the amount of information one variable provides about another, termed mutual information. In the intervening 75 years, this basic construction developed into a wide range of related quantities which aim to capture finer features of information. Unfortunately, there is no unique generalization of mutual information to the multivariate setting, though three stand out: total correlation (Watanabe, 1960), dual total correlation (Han, 1978) and interaction information (McGill, 1954). These approaches provide a holistic view of the amount of information that all sources under consideration provide about the target variable. Their key limitation, however, is that they do not tell you their relative importance.

Developing methods of “decomposing” multivariate information into such relative measures is an active area of research, first formalized by the “partial information decomposition” (PID) of William and Beer (Williams & Beer, 2010). The key concept that PID and subsequent decompositions aim to capture is that information from multiple sources can be broken down into pieces that arise uniquely, redundantly, and synergistically from combinations of those sources. In PID, each combination of sources and its associated contribution to the total correlation (Watanabe, 1960) is traditionally conceptualized via Venn-style diagrams (Fig. 1D) with each represented as a node in the lattice (Fig. 1E). To clarify the concepts of unique, redundant, and synergistic information, consider the “did it rain” example. In that case, an observation of current temperature on its own can be informative because rain tends to lower ambient temperature (unique information)—knowing the current temperature can help you guess whether or not it rained. If the temperature is high enough, you can be fairly certain that the water on the grass is not dew, making the dew point redundant. However, the dew point becomes relevant at lower temperatures such that temperature and dew point together (i.e., synergistically) provide information needed to rule out dew as an alternative explanation for the wet grass.

Applications of mutual information and its related information-theoretic quantities have been applied widely in biology (Farber et al., 1992; Fath et al., 2003; Hegyi & Garamszegi, 2011; Vicente et al., 2011), machine learning and data science (Huggins-Daines & Rudnicky, 2006; Shlezinger et al., 2020; Zhang et al., 2007), complex systems theory (Flecker et al., 2011; Lizier et al., 2010; Tehrani-Saleh & Adami, 2020), and geo-ecohydrology (Goodwell et al., 2020). This study is interested in whether information theoretic techniques can be used to learn about the evolutionary context of diverging populations. The premise is that the genome can be divided into protein coding (genic) and non-coding (intergenic) elements that evolve by different forces. Evolution of the coding regions is expected to be shaped by both the evolutionary characteristics of the species as well as natural selection with most loci under purifying selection and therefore showing reduced rate of accumulation of mutations among diverging lineages. Selection should have relatively little effect on non-coding regions.

Measures of genetic diversity and divergence are numerous and capture related but slightly different aspects of population evolution. The feature that some metrics (F_ST_ D_XY_) are expected to reflect more similar properties than others (D_XY_, π) depending on evolutionary history means their unique, redundant, and synergistic information can be exploited to reveal something about the context in which they evolved, for example whether they diverged with or without gene flow from a neighboring population. With this rationale, we computed eight common genetic metrics of simulated population genomic data under a suite of evolutionary scenarios. We chose coding/non-coding as the target variable and the value of each genetic metric in that region as a source variable and decomposed the multivariate mutual information over those sources using the PID of (Williams & Beer, 2010; see Figure 1). Using the architecture of the genome as the target variable (coding vs. noncoding), we believe this information signal is effectively standardized within a dataset, making comparison across species possible. We then asked whether the genomic decomposition gave an information signal about the context under which those data evolved, namely whether gene flow did or did not occur during divergence. Finally, we asked whether this signal was observable despite organismal variation: i.e., recombination rate, mutation rate, etc.

In brief, we found the decomposition of genomic information yields a rich information-space (7,648,800 nodes in this case), many of which are informative to the evolutionary context in which those sequences evolved. Nodes were able to detect presence/absence of gene flow better than current SFS-based methods. More importantly, if features of Earth’s surface prevent gene flow among some populations, this study suggests that information signature may be “received” and recorded in organisms’ DNA and can be measured using the decomposition of multivariate mutual information.

## METHODS

### Genetic simulations & evolutionary analysis

We simulated population divergence scenarios in SLiM v3.3.2 (Haller & Messer, 2019) using house scripts that automated the parameter generation, simulations, and genetic metric calculations with EggLib (Evolutionary Genetics and Genomics Library) v3.0.0b21 (De Mita & Siol, 2012). All simulations started with a random seed and initial genome sequence of 60,000 nucleotides divided into ten “coding” and ten “noncoding” regions of 3,000 nucleotides each to simulate a small eukaryotic genome. The “coding” and “noncoding” regions were composed of a random assortment of 1,000 codons (excluding stop codons) and 3,000 nucleotides, respectively. We simulated data under all combinations of the following four parameters: recombination rate r E {0, 10^-7^}, mutation rate µ E {2.5 x 10^-7^, 2.5 x 10^-6^}, symmetrical gene flow M E {0.00, 0.01, 0.10}, and effective population size N_e_ E {500, 1000} for a total of 24 scenarios (Table S1).

To simulate purifying selection on coding regions and calculate fitness, in SLiM we translated and compared the coding regions for each individual per generation to its initial (ancestral) amino acid sequence. The individual was assigned a reduced fitness of 0.9 based on one amino acid mismatch and 0.5 for two or more mismatches. Simulations were run for 100,000 generations and VCF (Variant Call Format) files were generated every 10,000 generations; each parameter combination (i.e. scenario) was replicated 100 times, yielding 24,000 VCF files that were then aggregated for genetic calculations (see Table S2). The pipeline to parallelize SLiM simulations for batch-running on high performance computing clusters and automate the processing of intermediate files and calculation of genetic metrics was written in Python, Jinja, Linux, and Eidos and is available at https://github.com/mmoral31/SLiM_pipeline.

To calculate genetic metrics, we used EggLib (Evolutionary Genetics and Genomics Library) on the aggregated VCFs generated by SLiM. The house script calculated eight statistics: F_ST_, D_a_, Tajima’s D, θ_W_, π, S, Fu’s FS, and D_XY_ in each 3-kb coding and noncoding region individually (i.e. window size and step size of 3 kb). Whether the region was “coding” or “noncoding” was recorded in the output for each window. These statistics were then passed into Imogen.jl for PID calculations (https://github.com/ELIFE-ASU/Imogen.jl).

### Technical variation & early divergence simulations

To assess technical variance of PID calculations, we generated an additional 300 genetic simulation replicates (for a total of N=400 per parameter combination) for a subset of parameters: M = 0, µ = 2.5 x 10^-7^, N_e_ E {500, 1000}, and r E {0, 10^-7^} (Table S2). To confirm the theoretical assumption that the genetic metrics hold no information at the immediate onset of divergence, and to test the reliability of decompositions during early-divergence, we ran another set of simulations focusing on early time-series sampling. For this we re-ran all 24 scenarios for 10,000 generations and sampled every 1,000 generations, with another 100 replicates per scenario (Table S2). The VCFs were generated at generation two instead of generation one due to software limitations. This set of data also consisted of 24,000 VCF files that were aggregated for genetic analysis.

### Partial information decomposition

#### Implementation & running of analyses

The preceding analysis provided a sequence of values for each genetic metric along with whether or not the associated 3-kb region was a coding or noncoding region. In other words, over the length of the genome, we had a binary vector c^➔^ representing whether or not a given region is coding, and eight vectors d^→^—one for each evolutionary statistic calculated in the region. We treated these as sequences of observations of random variables c and d_i_ which allowed us to ask how the information about whether a given region is coding decomposes over combinations of the evolutionary metrics using the PID formalism. However, in general the d^→^ are real-valued, and since the Williams and Beer PID formalism is limited to discrete-valued data, we performed a discretization process on each ^-^d**^-^**^➔^. For the sake of simplicity and to reduce systematic errors due to the short genome lengths (60,000 bp), we opted to use a “mean-threshold” binning scheme: the jth value of d^-**-**➔^, written here as d, was replaced with a 1 if it was greater than the average of all values in d^→^, and 0 otherwise. This yielded a set of eight binary vectors D^→^. PID could then be applied to the resulting vectors c^➔^ and D^→^_l_. In this process, the multivariate mutual information between c^➔^ and -_D_--➔_l_ was estimated and subsequently decomposed into a sum of non-negative terms, roughly describing the degree to which combinations of the variables D^→^_l_ provided unique, redundant, and synergistic information about whether or not a given region is coding. Due to computational limitations of PID, the process can only be applied to at most five of the evolutionary metrics at a time. In this work, we limited calculations to at most combinations of four metrics considered simultaneously to limit computation time and keep the amount of data generated manageable. We carried out the computation for all lattices with between one and four of the genetic metrics as source variables and the region classification c^➔^ as the target to yield 162 PID lattices per simulation per sampled generation for a total of 97,200 lattices containing 7,648,800 nodes in total.

#### Information clusters & informative metrics

For each node in each lattice per dataset, we created a vector of the partial information (n) for that node over time. Many of the resulting curves had similar temporal dynamics but started out with a different initial value. To account for this, we shifted each vector by subtracting the zeroth element, element-wise, to make all vectors start at the same zero value. To reduce the 764,880 vectors into classes of information signatures over time, we performed k-means clustering based on Euclidean norm distance in the 10-dimensional space formed by the 10 sampled timepoints. We used the “elbow method” to identify a reasonable number of clusters for the k-means analysis. That is, we plotted the explained variance against the number of clusters for a range of values of k, and selected k at or near a prominent elbow (bend) in the curve. Based on this method, we chose a 7-means clustering to reduce the number of patterns on which to conduct further analyses without oversimplifying the diversity of results.

#### Limitations

We note several aspects of this analysis that could affect the results obtained. First, the genomes simulated were small (60 kbp) relative to the gigabasepair-size genomes of vertebrates, for example. For simplicity, simulations did not incorporate standing variation in the population, which is a simplification of reality and could affect the rate at which nodes become informative. Further, technical aspects of the Williams and Beer formalization of PID, such as the lack of localizability, continuity and differentiability, leave room for further exploration. More recent work (Finn & Lizier, 2018; Makkeh et al., 2021) made progress in addressing these points, and have arguably developed notions of information decomposition that are easier to interpret (which may be important for the future of PID for biological applications). The choice to use Williams and Beer’s formalism necessitated a binning procedure on the computed evolutionary quantities before analysis, and the binning method can affect the resulting decompositions. It would be worth considering different discretization methods (i.e. Doane, 1976; Scargle et al., 2013; Scott, 1985) to assess the sensitivity of the results presented in this work.

#### Benchmarking against SFS simulations

To compare how well gPID recovered signatures of historical migration compared to a method based on site frequency spectrum analysis, we used the δaδi pipeline developed by Portik, et al. (Portik et al., 2017). Three technical replicates from each migration scenario were selected (M = 0, M = 0.01, and M = 0.1); all other parameters were held constant. Because the amount of population divergence after splitting increases with more generations, we used δaδi separately on data that were 10,000 generations and 100,000 generations post-split. Since the populations split at the same time and had constant population sizes post-split, we selected the divergence with no migration and divergence with symmetrical migration models for δaδi. The difference in these two models is one parameter (migration); thus, AIC scores should not be penalized from model complexity. Furthermore, it allowed us to test whether our varying migration scenarios would follow an expected trend: e.g., M = 0 data sets should have lower AIC values with the no migration model than the symmetrical migration model.

The merged VCF from each of the two populations from each migration scenario were converted into a folded allele frequency spectrum for δaδi to analyze. Two optimization methods were used to evaluate model fit (Nelder-Mead and BFGS; Gutenkunst et al., 2009) and AIC scores across technical replicates to determine the best model per migration scenario.

## RESULTS

Results yielded 97,200 PID lattices that contained 7,648,800 nodes in total. The long time series (10^5^ generations) showed that nodes within decomposed lattices exhibited partial information patterns that fell into seven timeseries classes (Figure 2B). Most nodes were invariable (N=302,207), while some reliably decreased (Class 2, N=196) or increased (Class 3, N=128). Complex time signatures are in SI (Figure S1).

**Figure 2.**
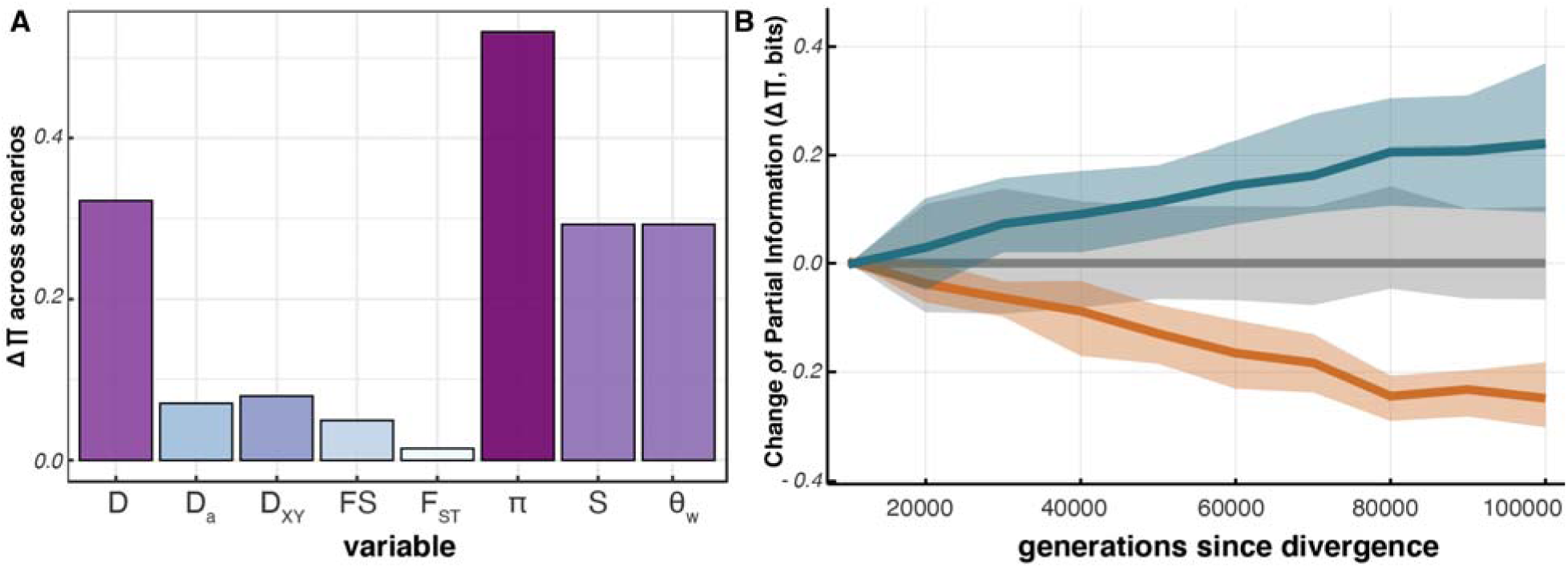
Summary of information content in genomic metrics. **(A)** Change in information content (Δ∏) in single-variable decompositions across evolutionary scenarios. A higher value reflects a greater ability for that variable to discriminate among evolutionary scenarios. **(B)** Time series summary of how nodes from the decomposed lattices change over time. Patterns were summarized into seven time-signature clusters using k-means. Three of those are shown here: flat (gray, N=303151), increasing (turquoise, N=368), and decreasing (orange, N=196).

Our exploration of evolutionary parameter space (*N_e_*, Μ, r, and μ) revealed the presence of gene flow was the easiest signal to detect (Figure 3), which is consistent with results based on analysis of the SFS (Gutenkunst et al., 2009). The variable most able to recover the gene flow signal was π while Tajima’s D and F_ST_ reflect the presence/absence of gene flow only at higher mutation rates (Figure S5). No nodes effectively discriminated between recombination rates, which could be because our metrics did not reflect patterns of genomic linkage. Results for technical variation showed the mean absolute distance (MAD) between any pair of lattices never exceeded *e*^-4^ bits, which is orders of magnitude below the differences observed between evolutionary scenarios.

**Figure 3.**
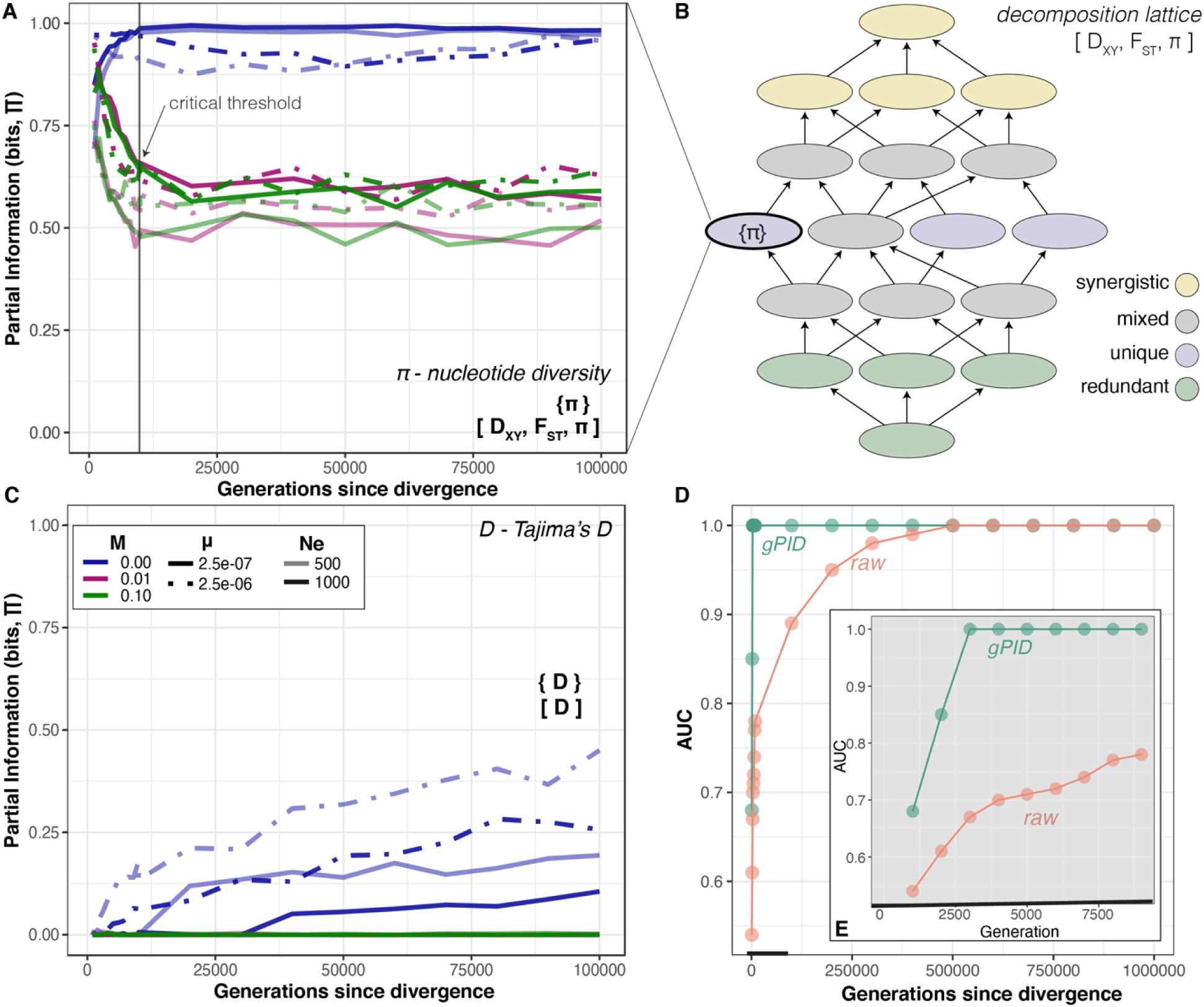
Discrimination of time-series signatures - (A) Partial information for Unique{Pi} from [D_XY_, F_ST_, π] over 100,000 generations, which shows the strongest discrimination capability between gene flow (pink, green) and no gene flow (blue) following a critical threshold when the pattern stabilizes. **(B)** The 18-node decomposition lattice for [D_XY_, F_ST_, π], from which a single node {π} is plotted in (a). (C) Single variable decomposition of Tajima’s D showing increasing partial information for {D} with time since divergence in the absence of gene flow, causing redistribution of information across lattices. **(D)** Comparison of AUC values for a receiver-operator curve (ROC) for the ability of π to discriminate between presence/absence of gene flow using gPID (green) versus raw π values (orange) over longer divergence (100,000 generations) and **(E)** early divergence (10,000 generations).

It is expected that before divergence begins that all nodes have negligible partial information because the simulated populations start out from a single sequence and that following divergence they rapidly departed from 0 bit. Results from the re-sampling at a higher frequency (every 10^3^ generations) are consistent with the assumption that the information content begins at 0 bit and becomes nonzero following the first generation.

Finally, we found that combinations of variables exhibited “additive” discriminatory behavior. As an example, the single variable decomposition [S] displayed sensitivity to both Μ (presence/absence) and μ (Figure S4A). The single variable decomposition of [D] reflected primarily presence or absence of gene flow (Μ, Figure 3C). The two-variable decomposition [S,D] presents a simple combination of [S] and [D], with a primary signal of gene flow (present in both [S] and [D]) and a secondary signal of μ (present in [S]).

## DISCUSSION

To understand the information signature of genetic metrics under different evolutionary settings, we simulated 60 kbp of “genic” and “intergenic” sequence data of individuals in each of two populations under 24 evolutionary scenarios. Evolutionary scenarios varied mutation rate (μ), recombination rate (r), effective population size (*N_e_*), and gene flow (Μ). We sampled genomes of 10 individuals per population at 10 timepoints per simulation to obtain a timeseries signature during divergence. We calculated eight genetic metrics (F_ST_, D_a_, Tajima’s D, θ_W_, π, S, Fu’s FS, and D_XY_) for each genomic element and decomposed the information signature of those using PID over lattices constructed from combinations of up to four variables using genic/intergenic as the target variable.

### The informative nodes

We find that information signatures depend almost entirely on the informativeness of a few nodes within the lattice representing key genetic metrics and not on the set of variables in the decomposition (Figures 2A, 3, S2, S3). The only exception to this pattern is Tajima’s D, which increases in informativeness over time in the absence of gene flow (Figure 3C). Consequently, when included among the variables Tajima’s D redistributes information over the lattice, making other nodes more informative even though Tajima’s D itself is rarely (if ever) part of the node with an ability to discriminate among evolutionary scenarios (e.g., Figure 3D). The variables lending greatest informativeness (i.e., ability to discriminate among evolutionary scenarios) were nucleotide diversity (π) and Watterson’s estimator (θ_W_). The types of nodes with greatest discrimination potential are either unique or redundant; synergistic nodes are never discriminatory among evolutionary scenarios (Figure 3). We interpret this to be due to the redundancy or nonindependence of the genetic metrics and not a behavior of PID itself. More complex evolutionary scenarios (gene flow plus differential adaptation) may wind up being reflected in synergistic nodes.

### Detection of gene flow

Our exploration of evolutionary parameter space (*N_e_*, Μ, r, and μ) reveals the presence/absence of gene flow to be easy to detect (Figure 3), which is consistent with results based on analysis of the SFS (Gutenkunst et al., 2009). The discrimination is binary rather than continuous, identifying the presence, not the degree, of gene flow in our simulations. However, sampling more rates of gene flow could further resolve this pattern. Our gPID had higher sensitivity to detecting gene flow than δaδi, which is based on the SFS. Both algorithms used in δaδi detected the higher gene flow and no gene flow but had mixed results at low levels of gene flow. In gPID, the variable most able to recover the gene flow signal is π while Tajima’s D and F_ST_ capture the presence/absence of gene flow only at higher mutation rates (Figure S5). Differences in mutation rate is recovered to a lesser extent than the presence of gene flow and is reflected best by Fu’s FS. Segregating sites (S) and Watterson’s estimator (θ_W_) reflect differences in mutation rate but are also influenced by presence/absence of gene flow (Figure S6). Variables S and θ_W_ yielded identical results (Figure S6); this is expected because θ_W_ is calculated by correcting S for the number of samples observed which was consistent across our simulations. This nonetheless demonstrates PID’s consistency to uncover equivalent patterns despite calculation over values of different scales. In situations with variable sample size, these metrics might produce different decompositions.

### Detection of recombination rates

No nodes effectively discriminated between recombination rates, which could be because our metrics were not calculated across linkage blocks and therefore did not reflect patterns of genomic linkage. We speculate that decomposing alternative genetic metrics could recover a difference in recombination rate and performing sliding window analyses across longer sequences could capture this. For differences in *N_e_* (Figure S7), the magnitude of partial information was slightly inflated under smaller effective population sizes (Figure 3). It is possible that additional targeted exploration of nodes with a focus on discriminating among *N_e_* could uncover a more useful decomposition that specifically reflects differences in relative *N_e_*.

### Technical variation

We assessed technical variance of PID through additional simulations (Table S2) and the technical variance among genetic simulations and genomic PIDs was exceedingly small. After as few as 10,000 generations, many PID nodes exhibited partial information near the theoretical maximum of 1 bit. For early divergence simulations, there is slightly less stability in the early generations evidenced by the greater number of time series clusters compared to the 10^5^-generation dataset (N=20 vs N=7 groups, respectively; Figure S1 vs. S8). With additional work, it might be possible to leverage this latency period to measure signals of very early divergence (≲ 1000 generations, Figure S9). Aside from slightly reduced power to discriminate among evolutionary scenarios in early generations, the information patterns stabilize once a critical threshold is met (Figure 3A). This pattern means PID can be reliable when applied to genomic datasets of wild populations in which population divergence is presumed to be ≥3000 generations. Yet, the discrimination of gene flow was easily detected by 1,000 generations past initial divergence (Figure 3E).

### General observations of gPID

Several broadscale observations can be made about both the behavior of information decomposition and its application to genetic metrics and evolution. First, it is surprising that F_ST_, which is one of the most broadly used and interpretable statistics in evolutionary biology, was the least informative when decomposed, while more fundamental measures of diversity (π, S or θ_W_) drove information patterns. Also, key variables determined information patterns rather than the combination of variables in the decomposition (i.e., the set of source variables). This finding is important because computing PID on many variables is costly and can be difficult to interpret. Results here suggest that small lattices with a few key variables are sufficient for many purposes, and new applications of PID should assess patterns within small decompositions first.

Likewise, we were surprised to find that the unique and redundant nodes far outweighed information in synergistic nodes. We interpret this to be a property of the genetic metrics analyzed, particularly their non-independence, and how the redundancy of the information they provide changes under different evolutionary settings. If, unlike other multivariate statistics, PID leverages interpretational power from the nonindependence of variables, that may be its biggest strength. It is possible that synergistic terms would gain power under more evolutionarily complex scenarios, such as reduced gene flow from a physical barrier in combination with differential adaptation, which remains a stumbling block for application of the SFS (Mathew & Jensen, 2015).

Finally, the decompositions show additive behavior among variables. For example, results from the two-variable decomposition [S, D] presents a simple combination of the individual patterns of [S] and [D]. Both [S] and [D] were able to detect gene flow, while [S] was secondarily able to detect differences in μ (Figures S4a, S4b). The joint decomposition of [S, D] primarily reflected presence/absence of gene flow (Μ, Figure 3C), with a secondary signal of μ, consistent with additive behavior (Figure S4C). If this is a true behavior of the method, then this should enable a targeted search of decomposition-space for lattices and nodes that will discriminate between parameters of interest. In theory, choice of metrics to decompose can be optimized based on which properties of a diverging population are of interest. Machine-learning approaches that can mine the large decomposition space for new useful metrics is a promising area for future work.

Using the techniques employed in this work, one could conceivably distill down diversity measures that specifically target particular aspects of speciation. For example, there exists, within the context of the [Dxy, Fst, π] lattice, a closed-form expression for {π}. If the work presented here translates to real-world, empirical data, then this closed form would represent a new genetic metric tailored specifically toward identifying gene flow between two diverging populations. Similar specialized analytics could, in principle, be derived for the other nodes or genetic metrics that prove consistent and useful.

### Is there a communication channel between Earth and life?

Results from this study demonstrate that the divergence of lineages can be measured using multivariate information of the genome at a population scale. While we did not measure the information content of the landscape, it is often supposed that the movement of individuals and their local adaptation is in part mediated by the texture and composition of Earth’s surface and climate. If that is true, then our simulations indicate those effects are communicated and recorded in evolutionary signatures of the genome. This message communicated from the landscape to lineage divergence, then, can be measured through the decomposition of mutual information. If every evolutionary lineage is a separate (though not phylogenetically independent) receiver of that message, it presents a simple conjecture: species that evolve strong population substructure in response to their landscape/environment receive that communication through a clear channel, while others, such as panmictic species or those whose population structure is less responsive to landscape features (e.g., whales or migrating birds) receive that communication through a noisy channel. The traits of the organism control the clarity of the channel. Assuming the communication is continuous over time and space, the generation time of the receiver (a lineage) would determine the temporal window at which the communication channel exists, while its geographic range determines its spatial footprint. In this study we did not measure the information embedded in the landscape. But, the message communicated by some regions of Earth, i.e., geological, climatic, or geographic settings, may be more complex, multifaceted, or information-rich than others. Whether these information-rich landscapes communicate that “message”, which we record as a signal of high diversification and/or biodiversity, is an open question.

## CONCLUSIONS

In summary, many outstanding questions in speciation and macroevolution would benefit from the ability to characterize aspects of lineage divergence across diverse organisms (Dolby et al., 2022). Gene flow is a simple example that is influenced by the geographical context of lineage divergence. We outline a new information theoretic approach to do this which leverages the architecture of the genome and partial redundancy of common genetic metrics. Application of PID here shows the presence/absence of gene flow is easily detectable by several decompositions despite differences in genomic or biological characteristics, such as recombination and mutation rates. PID of genomic information (gPID) is a promising and integrative analytical framework that can be applied to other variables and the large decomposition-space can be mined for nodes targeted to specific evolutionary characteristics and timepoints of interest. The application of communications theory in this context suggests the landscape or physical “context” is the sender of a communication which is received by lineages living on the landscape and encoded in the non-random structuring of genomic information among populations of each species. Those signatures can be measured using the partial decomposition of multivariate mutual information.

## ACKNOWLEDGEMENTS

We thank the Baja GeoGenomics consortium for useful feedback and discussions and Kenro Kusumi for support.

## DATA & CODE AVAILABILITY

Code is available on Github: https://github.com/elife-asu/Imogen.jl and https://github.com/mmoral31/SLiM_pipelineData files are available at https://www.dropbox.com/sh/8umfqmby1f0cmqj/AAC1EbdUfhw8XyyUGb4F5OAna?dl=0 and will be publicly archived upon publication.

## AUTHOR CONTRIBUTIONS

DGM and GAD conceived of this project; DGM, SIW, and GAD supervised analyses. MM carried out genetic simulations and pipeline automation, DGM performed information theoretic analyses. ARB performed site-frequency-spectrum analyses. GAD and DGM drafted and all authors critically revised the manuscript.

## REFERENCES

1. Antonelli, A. (2017). Biogeography: Drivers of bioregionalization. Nature Ecology & Evolution, 1(4), 0114. 10.1038/s41559-017-0114

2. Antonelli, A., Kissling, W. D., Flantua, S. G. A., Bermúdez, M. A., Mulch, A., Muellner-Riehl, A. N., Kreft, H., Linder, H. P., Badgley, C., Fjeldså, J., Fritz, S. A., Rahbek, C., Herman, F., Hooghiemstra, H., & Hoorn, C. (2018). Geological and climatic influences on mountain biodiversity. Nature Geoscience, 11(10), 718–725. 10.1038/s41561-018-0236-z

3. Araya-Donoso, R., Baty, S. M., Alonso-Alonso, P., Sanín, M. J., Wilder, B. T., Munguia-Vega, A., & Dolby, G. A. (2022). Implications of barrier ephemerality in geogenomic research. Journal of Biogeography, 49(11), 2050–2063. 10.1111/jbi.14487

4. Baker, P. A., Fritz, S. C., Dick, C. W., Eckert, A. J., Horton, B. K., Manzoni, S., Ribas, C. C., Garzione, C. N., & Battisti, D. S. (2014). The emerging field of geogenomics: Constraining geological problems with genetic data. Earth-Science Reviews, 135, 38–47. 10.1016/j.earscirev.2014.04.001

5. Chase, M. A., Stankowski, S., & Streisfeld, M. A. (2017). Genomewide variation provides insight into evolutionary relationships in a monkeyflower species complex ( *Mimulus* sect. Diplacus). American Journal of Botany, 104(10), 1510–1521. 10.3732/ajb.1700234

6. Crisci, J. V. (2001). The voice of historical biogeography. Journal of Biogeography, 28(2), 157–168. 10.1046/j.1365-2699.2001.00523.x

7. De Mita, S., & Siol, M. (2012). EggLib: Processing, analysis and simulation tools for population genetics and genomics. BMC Genetics, 13(1), 27. 10.1186/1471-2156-13-27

8. Doane, D. P. (1976). Aesthetic Frequency Classifications. The American Statistician, 30(4), 181–183.

9. Dolby, G. A. (2021). Towards a unified framework to study causality in Earth–life systems. Molecular Ecology, 30(22), 5628–5642. 10.1111/mec.16142

10. Dolby, G. A., Bennett, S. E. K., Dorsey, R. J., Stokes, M. F., Riddle, B. R., Lira-Noriega, A., Munguia-Vega, A., & Wilder, B. T. (2022). Integrating Earth–life systems: A geogenomic approach. Trends in Ecology & Evolution, 37(4), 371–384. 10.1016/j.tree.2021.12.004

11. Farber, R., Lapedes, A., & Sirotkin, K. (1992). Determination of eukaryotic protein coding regions using neural networks and information theory. Journal of Molecular Biology, 226(2), 471–479. 10.1016/0022-2836(92)90961-I

12. Fath, B. D., Cabezas, H., & Pawlowski, C. W. (2003). Regime changes in ecological systems: An information theory approach. Journal of Theoretical Biology, 222(4), 517–530. 10.1016/S0022-5193(03)00067-5

13. Finn, C., & Lizier, J. (2018). Pointwise Partial Information Decomposition Using the Specificity and Ambiguity Lattices. Entropy, 20(4), 297. 10.3390/e20040297

13a. Flecker, B., Alford, W., Beggs, J. M., Williams, P. L., & Beer, R. D. (2011). Partial information decomposition as a spatiotemporal filter. Chaos: An Interdisciplinary Journal of Nonlinear Science, 21(3), 037104. 10.1063/1.3638449

14. Goodwell, A. E., Jiang, P., Ruddell, B. L., & Kumar, P. (2020). Debates—Does Information Theory Provide a New Paradigm for Earth Science? Causality, Interaction, and Feedback. Water Resources Research, 56(2). 10.1029/2019WR024940

15. Gutenkunst, R. N., Hernandez, R. D., Williamson, S. H., & Bustamante, C. D. (2009). Inferring the Joint Demographic History of Multiple Populations from Multidimensional SNP Frequency Data. PLoS Genetics, 5(10), e1000695. 10.1371/journal.pgen.1000695

16. Haller, B. C., & Messer, P. W. (2019). SLiM 3: Forward Genetic Simulations Beyond the Wright– Fisher Model. Molecular Biology and Evolution, 36(3), 632–637. 10.1093/molbev/msy228

17. Han, T. S. (1978). Nonnegative entropy measures of multivariate symmetric correlations. Information and Control, 36(2), 133–156. 10.1016/S0019-9958(78)90275-9

18. Hegyi, G., & Garamszegi, L. Z. (2011). Using information theory as a substitute for stepwise regression in ecology and behavior. Behavioral Ecology and Sociobiology, 65(1), 69–76. 10.1007/s00265-010-1036-7

19. Hendry, A. P., Bolnick, D. I., Berner, D., & Peichel, C. L. (2009). Along the speciation continuum in sticklebacks. Journal of Fish Biology, 75(8), 2000–2036. 10.1111/j.1095-8649.2009.02419.x

20. Heslop-Harrison, J. S. (Pat), Schwarzacher, T., & Liu, Q. (2023). Polyploidy: Its consequences and enabling role in plant diversification and evolution. Annals of Botany, 131(1), 1–10. 10.1093/aob/mcac132

21. Huggins-Daines, D., & Rudnicky, A. I. (2006). A constrained baum-welch algorithm for improved phoneme segmentation and efficient training. Interspeech 2006, paper 1580-Tue3A1O.2-0. 10.21437/Interspeech.2006-364

22. Liu, S., Zhang, L., Sang, Y., Lai, Q., Zhang, X., Jia, C., Long, Z., Wu, J., Ma, T., Mao, K., Street, N. R., Ingvarsson, P. K., Liu, J., & Wang, J. (2022). Demographic History and Natural Selection Shape Patterns of Deleterious Mutation Load and Barriers to Introgression across *Populus* Genome. Molecular Biology and Evolution, 39(2), msac008. 10.1093/molbev/msac008

23. Lizier, J. T., Prokopenko, M., & Zomaya, A. Y. (2010). Information modification and particle collisions in distributed computation. Chaos: An Interdisciplinary Journal of Nonlinear Science, 20(3), 037109. 10.1063/1.3486801

24. Makkeh, A., Gutknecht, A. J., & Wibral, M. (2021). Introducing a differentiable measure of pointwise shared information. Physical Review E, 103(3), 032149. 10.1103/PhysRevE.103.032149

25. Marques, D. A., Lucek, K., Haesler, M. P., Feller, A. F., Meier, J. I., Wagner, C. E., Excoffier, L., & Seehausen, O. (2017). Genomic landscape of early ecological speciation initiated by selection on nuptial colour. Molecular Ecology, 26(1), 7–24. 10.1111/mec.13774

26. Martin, S. H., Dasmahapatra, K. K., Nadeau, N. J., Salazar, C., Walters, J. R., Simpson, F., Blaxter, M., Manica, A., Mallet, J., & Jiggins, C. D. (2013). Genome-wide evidence for speciation with gene flow in *Heliconius* butterflies. Genome Research, 23(11), 1817– 1828. 10.1101/gr.159426.113

27. Mathew, L. A., & Jensen, J. D. (2015). Evaluating the ability of the pairwise joint site frequency spectrum to co-estimate selection and demography. Frontiers in Genetics, 6. 10.3389/fgene.2015.00268

28. McGill, W. (1954). Multivariate information transmission. Transactions of the IRE Professional Group on Information Theory, 4(4), 93–111. 10.1109/TIT.1954.1057469

29. Merrill, R. M., Dasmahapatra, K. K., Davey, J. W., Dell’Aglio, D. D., Hanly, J. J., Huber, B., Jiggins, C. D., Joron, M., Kozak, K. M., Llaurens, V., Martin, S. H., Montgomery, S. H., Morris, J., Nadeau, N. J., Pinharanda, A. L., Rosser, N., Thompson, M. J., Vanjari, S., Wallbank, R. W. R., & Yu, Q. (2015). The diversification of *Heliconius* butterflies: What have we learned in 150 years? Journal of Evolutionary Biology, 28(8), 1417–1438. 10.1111/jeb.12672

30. Myers, S., Fefferman, C., & Patterson, N. (2008). Can one learn history from the allelic spectrum? Theoretical Population Biology, 73(3), 342–348. 10.1016/j.tpb.2008.01.001

31. Naka, L. N., Werneck, F. P., Rosser, N., Pil, M. W., & Boubli, J. P. (2022). Editorial: The role of rivers in the origins, evolution, adaptation, and distribution of biodiversity. *Frontiers in Ecology and Evolution*, *10*, 1035859. 10.3389/fevo.2022.1035859

32. Portik, D. M., Leaché, A. D., Rivera, D., Barej, M. F., Burger, M., Hirschfeld, M., Rödel, M., Blackburn, D. C., & Fujita, M. K. (2017). Evaluating mechanisms of diversification in a Guineo-Congolian tropical forest frog using demographic model selection. Molecular Ecology, 26(19), 5245–5263. 10.1111/mec.14266

33. Rahbek, C., Borregaard, M. K., Antonelli, A., Colwell, R. K., Holt, B. G., Nogues-Bravo, D., Rasmussen, C. M. Ø., Richardson, K., Rosing, M. T., Whittaker, R. J., & Fjeldså, J. (2019). Building mountain biodiversity: Geological and evolutionary processes. Science, 365(6458), 1114–1119. 10.1126/science.aax0151

34. Ravinet, M., Yoshida, K., Shigenobu, S., Toyoda, A., Fujiyama, A., & Kitano, J. (2018). The genomic landscape at a late stage of stickleback speciation: High genomic divergence interspersed by small localized regions of introgression. PLOS Genetics, 14(5), e1007358. 10.1371/journal.pgen.1007358

35. Ribas, C. C., Fritz, S. C., & Baker, P. A. (2022). The challenges and potential of geogenomics for biogeography and conservation in Amazonia. Journal of Biogeography, 49(10), 1839–1847. 10.1111/jbi.14452

36. Rougemont, Q., Huber, B., Martin, S., Whibley, A., Estrada, C., Solano, D., Orpet, R., McMillan, W. O., Frérot, B., & Joron, M. (2023). Subtle Introgression Footprints at the End of the Speciation Continuum in a Clade of Heliconius Butterflies. Molecular Biology and Evolution, 40(7), 1–21. 10.1093/molbev/msad166

37. Samuk, K., Owens, G. L., Delmore, K. E., Miller, S. E., Rennison, D. J., & Schluter, D. (2017). Gene flow and selection interact to promote adaptive divergence in regions of low recombination. *Molecular Ecology*, *26*(17), 4378–4390. 10.1111/mec.14226

38. Scargle, J. D., Norris, J. P., Jackson, B., & Chiang, J. (2013). STUDIES IN ASTRONOMICAL TIME SERIES ANALYSIS. VI. BAYESIAN BLOCK REPRESENTATIONS. The Astrophysical Journal, 764(2), 167. 10.1088/0004-637X/764/2/167

39. Scott, D. W. (1985). Averaged Shifted Histograms: Effective Nonparametric Density Estimators in Several Dimensions. The Annals of Statistics, 13(3). 10.1214/aos/1176349654

40. Shannon, C. E. (1948). A Mathematical Theory of Communication. *Bell Systems Technical Journal*, *27*, 379–423.

41. Shlezinger, N., Farsad, N., Eldar, Y. C., & Goldsmith, A. J. (2020). ViterbiNet: A Deep Learning Based Viterbi Algorithm for Symbol Detection. IEEE Transactions on Wireless Communications, 19(5), 3319–3331. 10.1109/TWC.2020.2972352

42. Stankowski, S., Chase, M. A., Fuiten, A. M., Rodrigues, M. F., Ralph, P. L., & Streisfeld, M. A. (2019). Widespread selection and gene flow shape the genomic landscape during a radiation of monkeyflowers. PLOS Biology, 17(7), e3000391. 10.1371/journal.pbio.3000391

43. Stankowski, S., Chase, M. A., McIntosh, H., & Streisfeld, M. A. (2023). Integrating top-down and bottom-up approaches to understand the genetic architecture of speciation across a monkeyflower hybrid zone. Molecular Ecology, 32(8), 2041–2054. 10.1111/mec.16849

44. Supple, M. A., Papa, R., Hines, H. M., McMillan, W. O., & Counterman, B. A. (2015). Divergence with gene flow across a speciation continuum of Heliconius butterflies. *BMC Evolutionary Biology*, *15*(1), 204. 10.1186/s12862-015-0486-y

45. Tehrani-Saleh, A., & Adami, C. (2020). Can Transfer Entropy Infer Information Flow in Neuronal Circuits for Cognitive Processing? Entropy, 22(4), 385. 10.3390/e22040385

46. Vicente, R., Wibral, M., Lindner, M., & Pipa, G. (2011). Transfer entropy—A model-free measure of effective connectivity for the neurosciences. Journal of Computational Neuroscience, 30(1), 45–67. 10.1007/s10827-010-0262-3

47. Watanabe, S. (1960). Information Theoretical Analysis of Multivariate Correlation. IBM Journal of Research and Development, 4(1), 66–82. 10.1147/rd.41.0066

48. Williams, P. L., & Beer, R. D. (2010). *Nonnegative Decomposition of Multivariate Information* (arXiv:1004.2515). arXiv. http://arxiv.org/abs/1004.2515

49. Xue, A. T., & Hickerson, M. J. (2015). The aggregate site frequency spectrum for comparative population genomic inference. Molecular Ecology, 24(24), 6223–6240. 10.1111/mec.13447

50. Zhang, X., Wang, Y., & Zhao, Z. (2007). A Hybrid Speech Recognition Training Method for HMM Based on Genetic Algorithm and Baum Welch Algorithm. *Second International Conference on Innovative Computing*, Informatio and Control (ICICIC 2007), 572–572. 10.1109/ICICIC.2007.33

