## Supporting Information for "The information signature of diverging lineages"

**Table of Contents**

Table S1	Summary of parameter values for simulations

Table S2	Summary of number of simulations including technical/early simulations per variable combination

Table S3	Genetic metrics used in this study

Figure S1	Information clusters over time 100k gens

Figure S2	Nodes with decreasing information content over time

Figure S3	Nodes with increasing information content over time

Figure S4	Additivity behavior of nodes

Figure S5	Information detection with {F_ST_}

Figure S6	Information results of S vs. $\theta_{W}$

Figure S7	S and/or $\theta_{W}$ results with Dxy

Figure S8	Information clusters over time 10k gens

Figure S9	Early divergence & critical thresholds

Figure S10	Decompositions of pi and their time signatures

Figure S11	18-node lattice for 3-variable decomposition

Figure S12	Receiver operator curves for Ne, M, *µ*

Figure S13	Discrimination capability of gPID{π} versus raw π values

**Supporting Figures & Tables**

**Table 1 Summary of parameter values** used over genetic simulations in the program SLiM. In these simulations, gene flow/migration is defined as the probability that any given offspring will come from a defined source population. In total, there were 24 evolutionary scenarios simulated across the four variables. Here we define “scenario” as a unique combination of parameter values for which genetic data was simulated.

| **Variable** | **Symbol** | **Parameter values** |
| --- | --- | --- |
| Mutation rate | *µ* | $\in\left\{ 2.5\times{10}^{-7},2.5\times{10}^{-6} \right\}$ |
| Recombination rate | *r* | $\in\left\{ 0,{10}^{-7} \right\}$ |
| Effective population size | *Ne* | $\in\left\{ 500, 1000 \right\}$ |
| Gene flow/migration | *M* | $\in\left\{ 0.0, 0.01, 0.1 \right\}$ |

**Table 2 Summary of the number of simulated datasets** for the long time series, short time series, and PID technical replicates. For each dataset, genetic metrics were calculated per genic/intergenic region and then the replicates per scenario (^†^) were concatenated prior to analysis in PID, meaning that technical variance among genetic simulations is accounted for in the PID results. To assess technical variance in the calculation of the partial information *per se*, PID technical replicates (row three) were concatenated into batches of 100 (N=3) per scenario and then run through the PID analysis. PID results were then analyzed with PID results from corresponding datasets from the long time series (total N=4 datasets per scenario) and the mean-absolute-distance (MAD) was calculated to determine technical variance in the PID calculations, which we found to be negligible ($\leq e^{-4}$ bit).

| **Type** | **Number of generations simulated** | **Sampling interval (generations)** | **Number of datasets in time series** | **Number of scenarios simulated** | **Replicates per scenario**^†^ | **Total simulated datasets** |
| --- | --- | --- | --- | --- | --- | --- |
| *Long time series* | 100,000 | 10,000 | 10 | 24 | 100 | **24,000** |
| *Short time series* | 10,000 | 1,000 | 10 | 24 | 100 | **24,000** |
| *PID technical replicates* | 100,000 | 10,000 | 10 | 4 | 300 | **12,000** |

**Table 3** Summary of genetic metrics used in this study and their definitions and sources. These metrics characterize the genetic diversity and/or divergence of individuals or populations and were calculated in EggLib. ^1^Holsinger & Weir 2009, ^2^Cruickshank & Hahn 2014, ^3^Watterson 1975, ^4^Rédei 2008, ^5^RoyChoudhury & Wakeley 2010, ^6^Tajima 1989a, ^7^Tajima 1989b, ^8^Garrigan et al. 2009, ^9^Fu 1997.

| **Genetic metric** | **Name** | **Equation** | **Interpretation** |
| --- | --- | --- | --- |
| F_ST_ | Fixation index | $F_{st}=\frac{\left( p_{1}-p_{2} \right)^{2}}{2\bar{p}\left( 1-\bar{p} \right)}$ | Measures relative population differentiation normalized by intrapopulation diversity^1^ |
| D_XY_ | Absolute  differentiation | $D_{xy}=\sum_{ij} x_{i}y_{j}d_{ij}$ | Measures absolute population differentiation invariant to intrapopulation diversity^2^ |
| S | Segregating  sites | *n/a* | Measures sequence polymorphism as the number of site differences in a sequence alignment^3^ |
| π | Nucleotide  diversity | $\sum_{ij} x_{i}x_{j}\pi_{ij}$ | Quantifies genetic diversity as the average number of nucleotide differences between populations^4^ |
| θ_w_ | Watterson estimator | θ_w_ =$\frac{S}{\sum_{i=1}^{n-1} \frac{1}{i}}$ | Measure of genetic diversity based on normalization of the number of polymorphic sites^5^ |
| D | Tajima’s D | $D=\frac{\theta_{T}-\theta_{W}}{\sqrt{V\left( \theta_{T}-\theta_{W} \right)}}$ | Tests for selective neutrality but is sensitive to demographic effects^6,7,8^ |
| FS | Fu’s FS | $FS=\ln\left( \frac{\hat{S}}{1-\hat{S}} \right)$ | Tests for selective neutrality but is sensitive to demographic effects^8,9^ |
| D_a_ | Pairwise differentiation | $D_{a}=D_{xy}-\frac{1}{2}\left( \pi_{x}+\pi_{y} \right)$ | Nucleotide differences between populations since splitting, sensitive to unequal population variation^2^ |

**Supporting References**

Cruickshank TE, Hahn MW (2014) Reanalysis suggests that genomic islands of speciation are due to reduced diversity, not reduced gene flow. *Molecular Ecology*, **23**, 3133–3157.

Fu Y-X (1997) Statistical Tests of Neutrality of Mutations Against Population Growth, Hitchhiking and Background Selection. *Genetics*, **147**, 915.

Garrigan D, Lewontin R, Wakeley J (2009) Measuring the Sensitivity of Single-locus “Neutrality Tests” Using a Direct Perturbation Approach. *Molecular Biology and Evolution*, **27**, 73–89.

Holsinger KE, Weir BS (2009) Fundamental concepts in genetics: Genetics in geographically structured populations: defining, estimating and interpreting FST. *Nature Publishing Group*, **10**, 639–650.

Rédei GP (2008) *Encyclopedia of Genetics, Genomics, Proteomics, and Informatics*. Springer, Dordrecht.

RoyChoudhury A, Wakeley J (2010) Sufficiency of the number of segregating sites in the limit under finite-sites mutation. *Theoretical Population Biology*, **78**, 118–122.

Tajima F (1989a) Statistical method for testing the neutral mutation hypothesis by DNA polymorphism. *Genetics*, **123**, 585–595.

Tajima F (1989b) The effect of change in population size on DNA polymorphism. *Genetics*, **123**, 597–601.

Watterson GA (1975) On the number of segregating sites in genetical models without recombination. *Theoretical Population Biology*, **7**, 256–276.

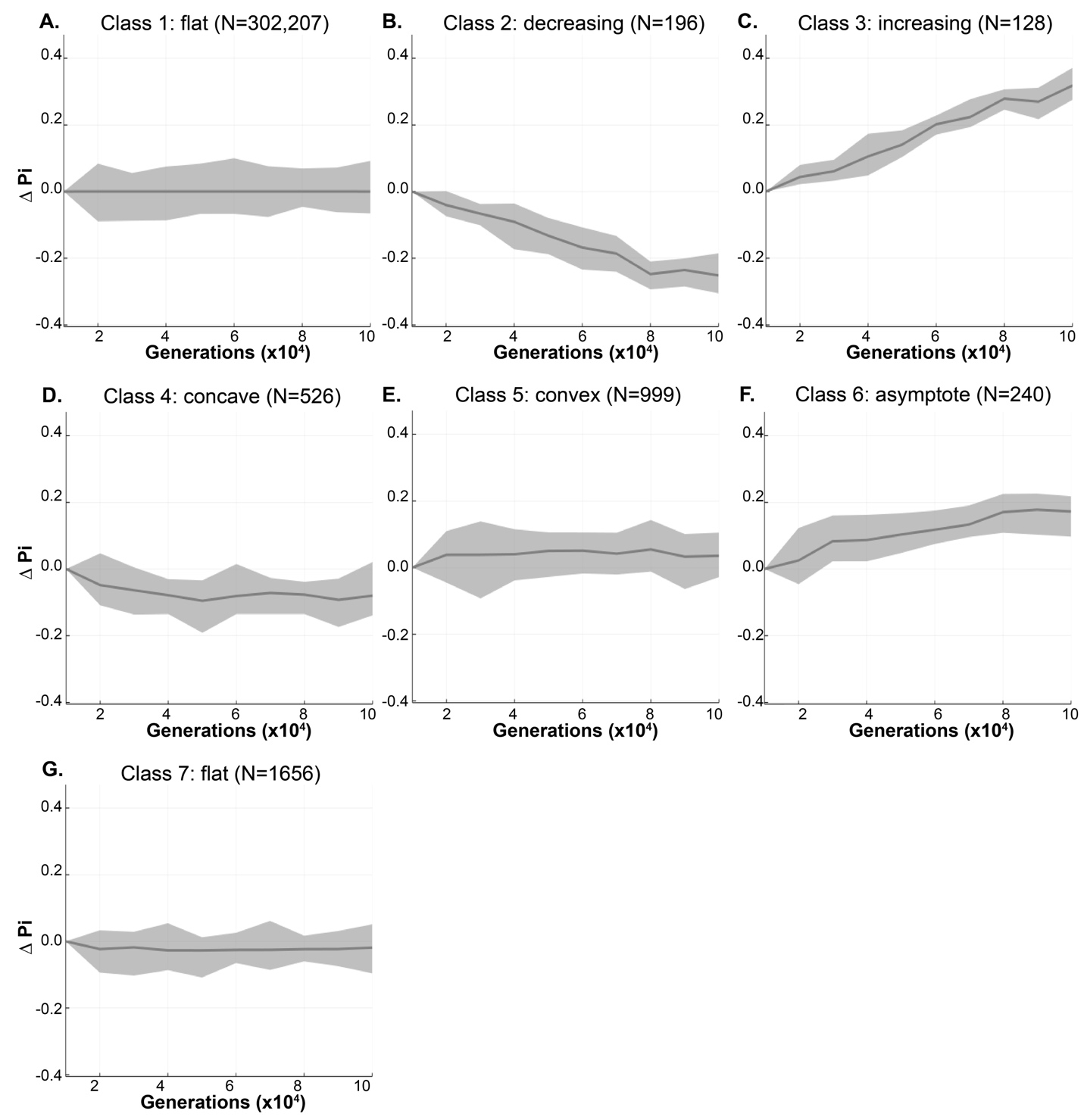

**Figure S1	Information signatures over time** clustered using K-means which resulted in seven total patterns (i.e. classes). These patterns resulted from nodes that were vectorized over sampling duration (sampled every 10,000 generations over 100,000 generations) generated from lattices based on 24 evolutionary scenarios (Table 1). Patterns are: A) uninformative/flat, B) decreasing informativeness over time, C) increasing informativeness over time, D and E) varying informativeness over time, F) increasing informativeness with asymptote, and G) a flat curve similar to class 1 but with more variance. The number of vectorized nodes (N) falling into each class are given in parentheses along with 100% envelope (grey). Each lattice is estimated based on 100 simulations.

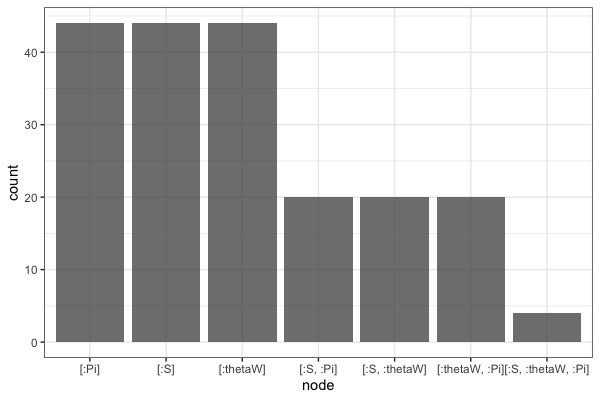

**Figure S2**	**Nodes with decreasing information over time** (from Figure 1, class two, N = 196). Single variables represent unique information, non-single variables represent redundant information amongst those variables. Single-variable nodes occur in more lattices than non-single variables, and therefore have higher counts. Variables are: D - Tajima’s D, thetaW - Watterson’s estimator, Pi - nucleotide diversity, S - segregating sites.

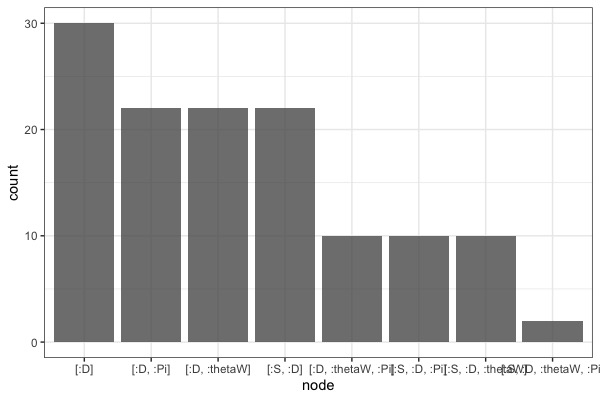

**Figure S3	Nodes with increasing information over time** (from Figure 1, class three, N=128). Single variables represent unique information, non-single variables represent redundant information amongst those variables. Single unique metrics occur in more decomposition lattices than non-single variables, and therefore have higher counts. Variables are: D - Tajima’s D, thetaW - Watterson’s estimator, Pi - nucleotide diversity, S - segregating sites.

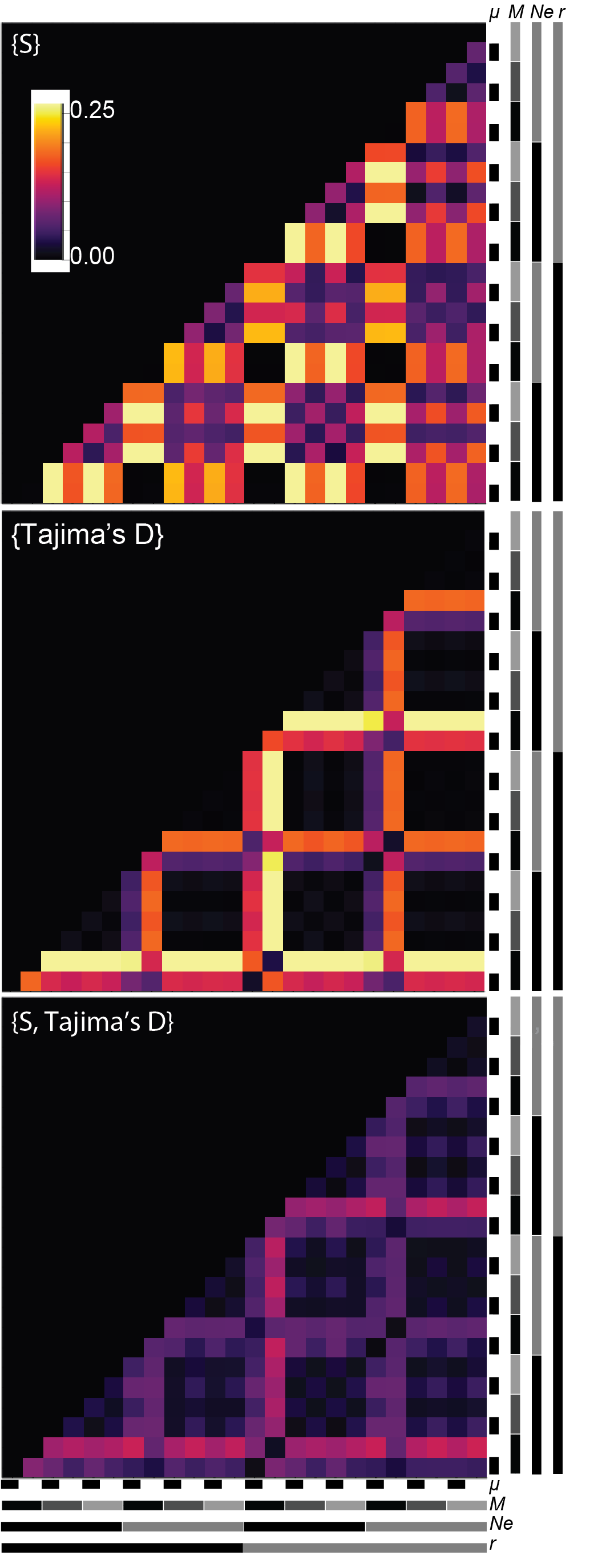

**Figure S4.** **Additivity behavior of nodes in decomposed lattices.** **(A)** Information variability across evolutionary simulations for single decomposition of segregating sites (S) and **(B)** Tajima’s D. Higher values (yellow) indicate stronger discrimination ability. Grey bars at bottom and right are guides for the heatmap pattern expected for each variable simulated (μ - mutation rate, M - gene flow, *N_e_* - effective population size, r - recombination rate). For example, Tajima’s D is discriminatory among gene flow whereas S detects both gene flow and mutation rate. **(C)** The information signatures are additive, wherein {S, Tajima’s D} displays the additive effects from panels A and B and the signal in heatmap **(C)** is exactly 1/4 the sum of the signals in **(A)** and **(B)**, demonstrating true additivity.

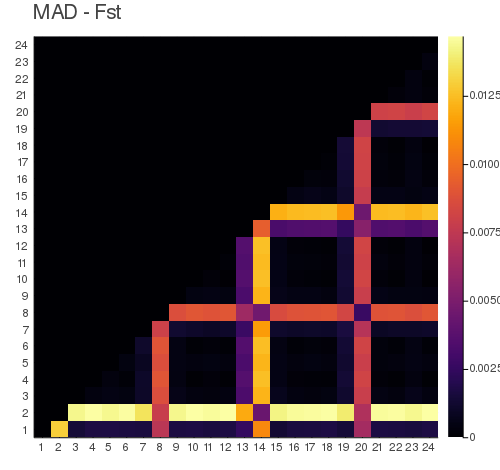

**Figure S5**	Heatmap showing differential information across evolutionary scenario based on Mean Absolute Distance (MAD) for F_ST_ as a single variable decomposition. Scale refers to bits of information in this node between scenarios with yellow being highest. The grey guides show the patterns that are expected depending on which of the evolutionary parameters is detectable with this node. The presence/absence of gene flow (light grey vs. dark grey and black for the M row) is the only pattern detected here.

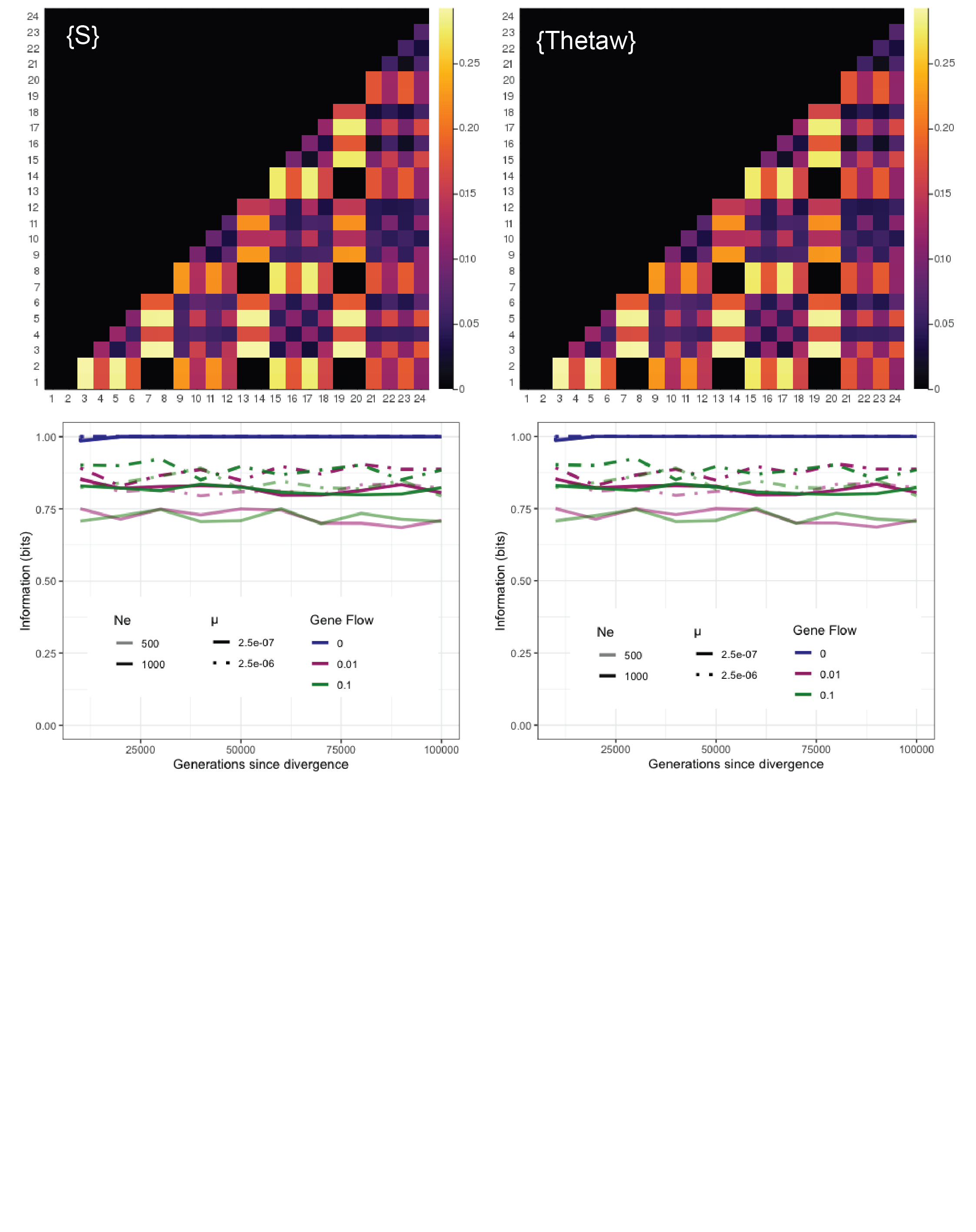

**Figure S6	Comparison of results from single decompositions of {S} (left) and {thetaw} (right).** Results are equivalent for the ability to discriminate amongst evolution scenarios (heatmaps, top) as well as information signatures over time under evolutionary scenarios modified by effective population size (Ne), mutation rate (μ) and gene flow. The pattern guides for the heatmaps are shown in Figures 4 and S6. This figure shows that although S and ThetaW have a different range of values, their equivalency is picked up by PID (thetaW is a normalized version of S).

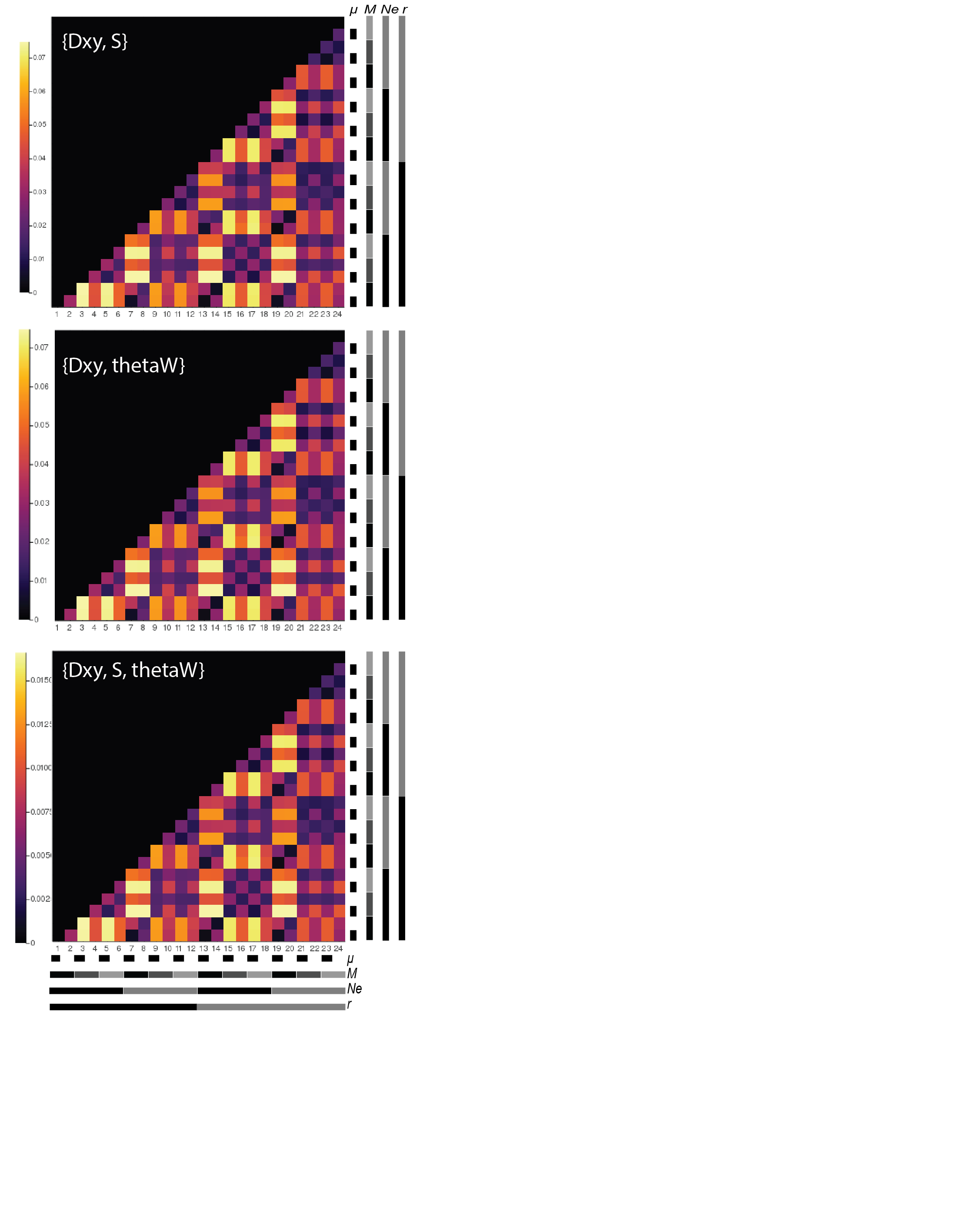

**Figure S7**	**S (top) and thetaW (middle) show equivalent patterns in the presence of additional variables (D_XY_)** and show a signal of both mutation rate and presence/absence of migration. The information content (scale bar to left) is decreased proportionally in the three-variable decomposition (bottom) because the mean absolute distance (MAD) is calculated over the larger matrix.

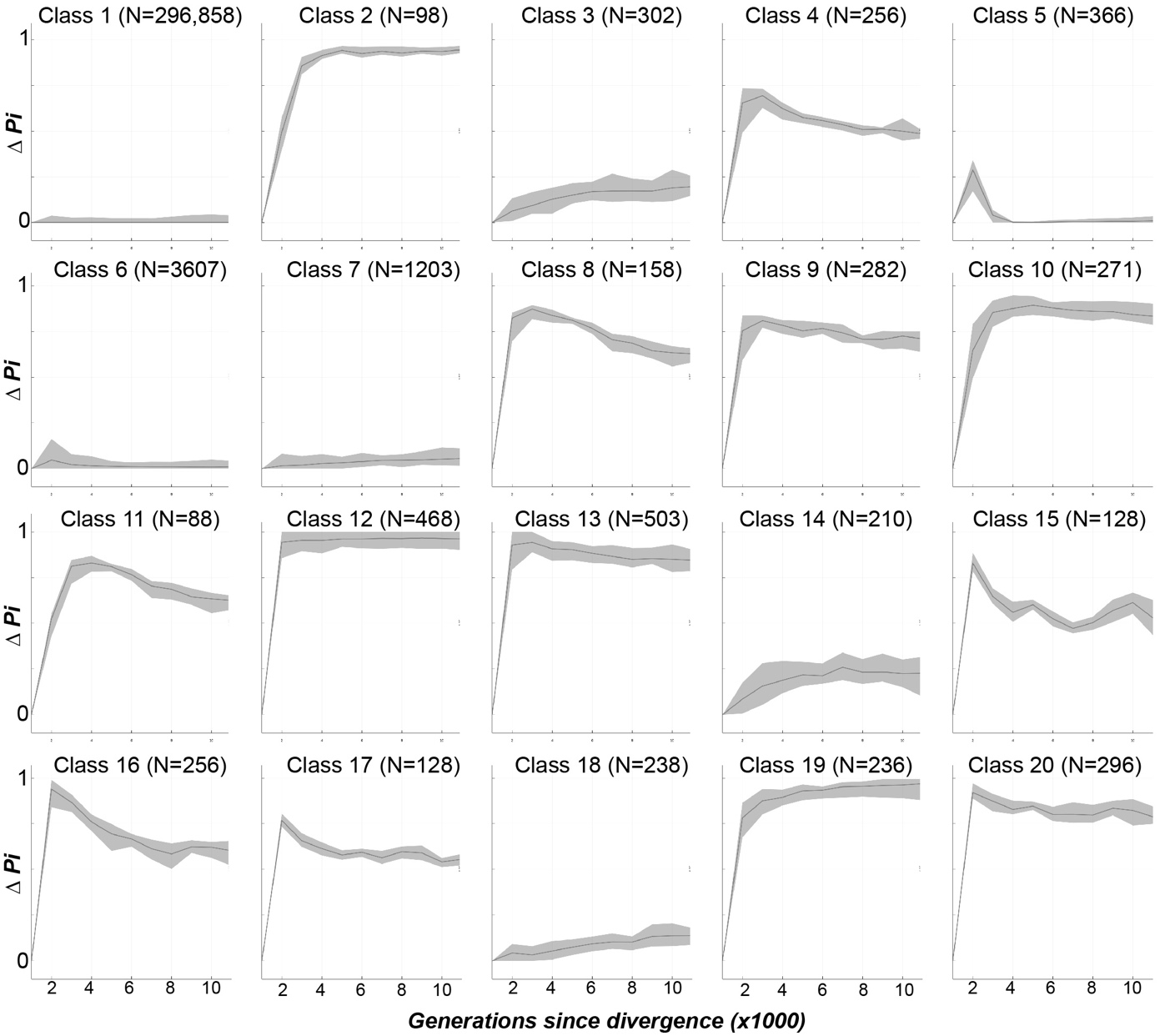

**Figure S8**	**Number of classes revealed by k-means clustering** in the short (10^4^ generation) dataset. There are more clusters (N=20) than observed for the longer generations (N=7, Figures 1, S1), which reflects the variable nature these metrics during early divergence. Note, all axes in this figure are identical.

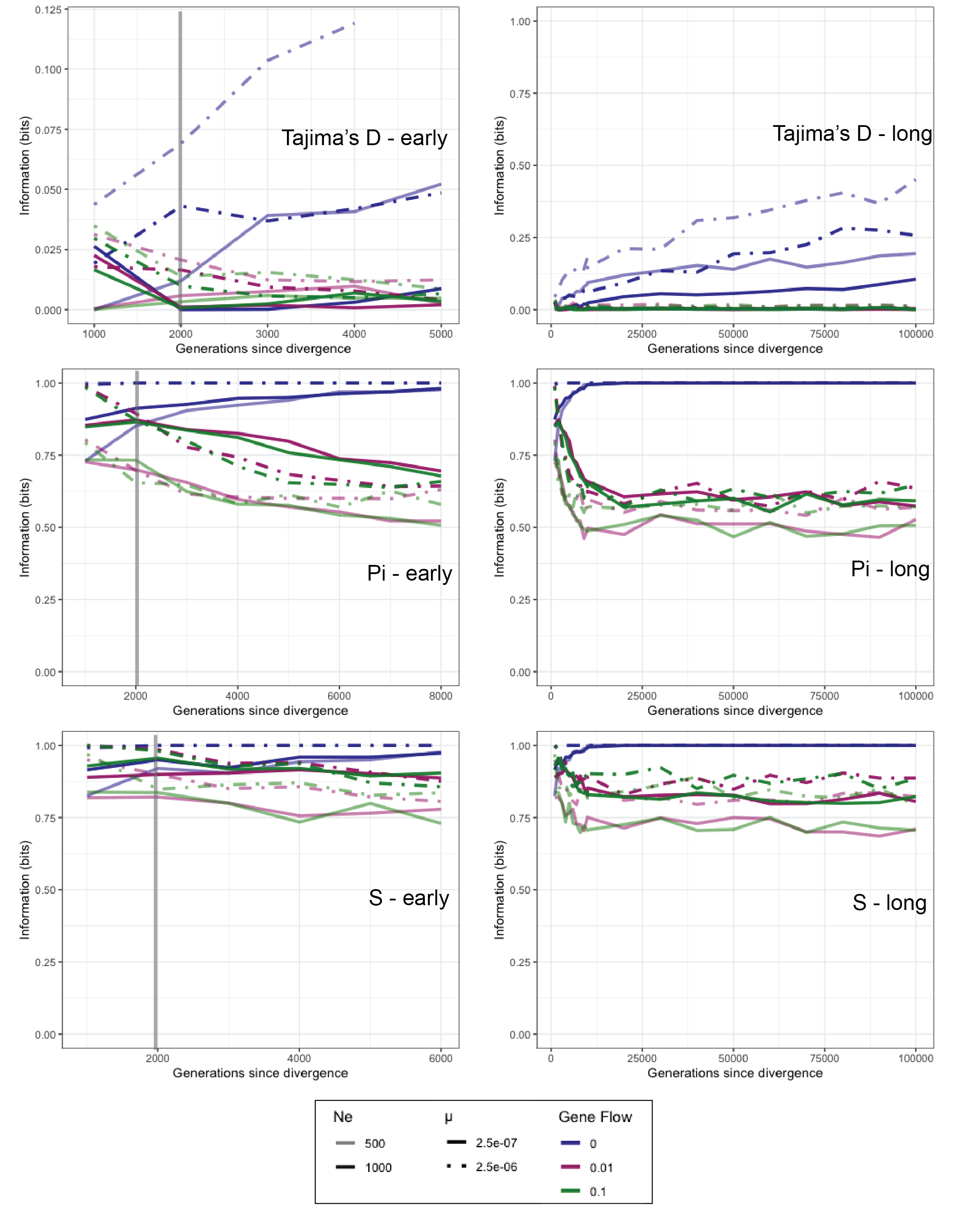

**Figure S9**	**Comparison of very early divergence relative to longer divergence** for three single-variable decompositions (Tajima’s D, Pi, and S). Vertical line denotes the 2000 generation mark, which in most simulations was a critical threshold at which the direction of the function (increasing or decreasing) is generally stable. All decomposed nodes have a theoretical start point of 0 bit, after which the departure from zero is rapid if the node is informative. Following a critical threshold, the node’s information stabilizes.

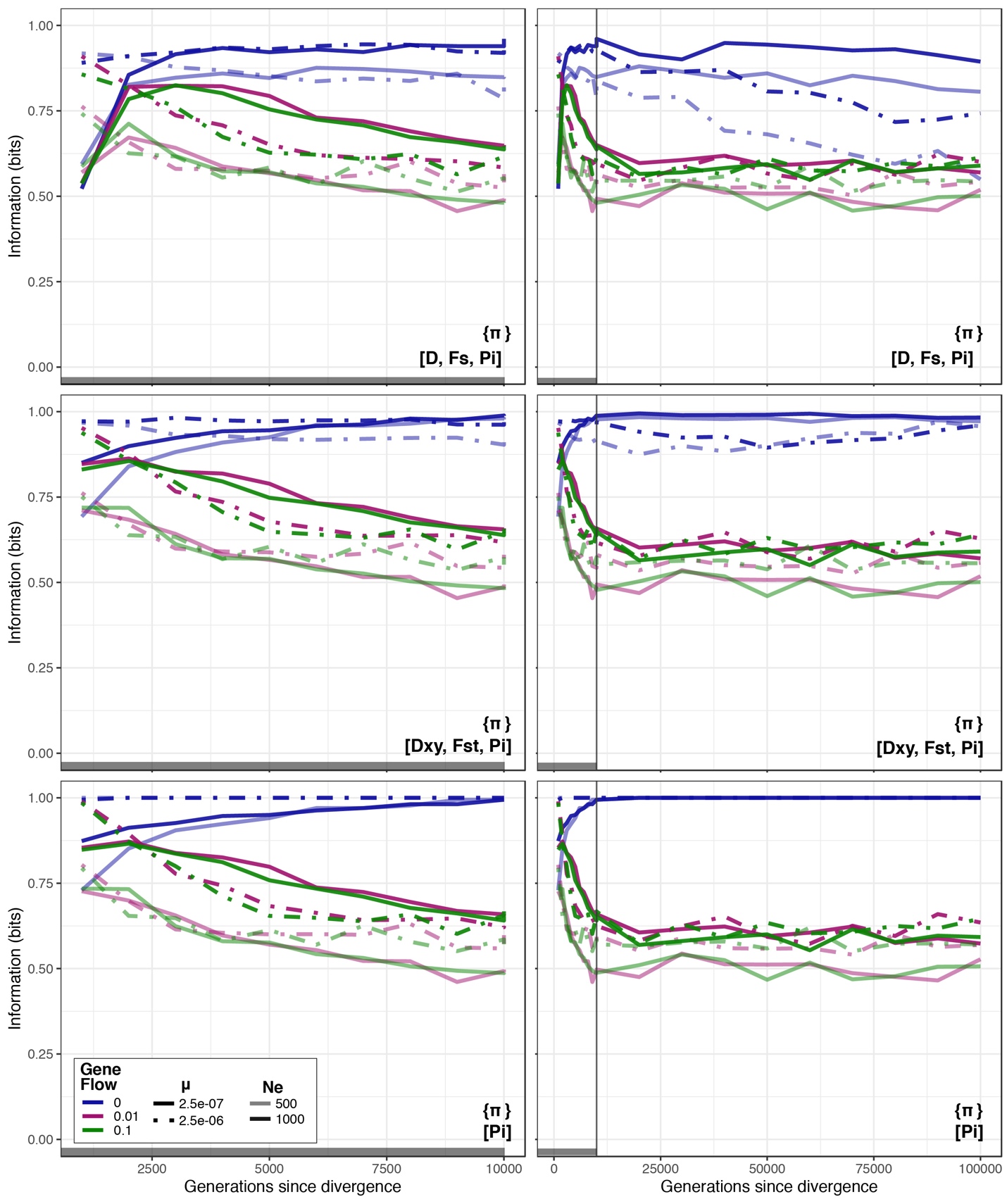

**Figure S10	Information content of unique(Pi) over time can discriminate between absence (blue) and presence (pink, green) of gene flow**. Its informativeness depends somewhat by the presence of perturbing variables (Tajima’s D, top), but not Dxy or Fst (middle). The early divergence signal (left) and long divergence signals are both shown.

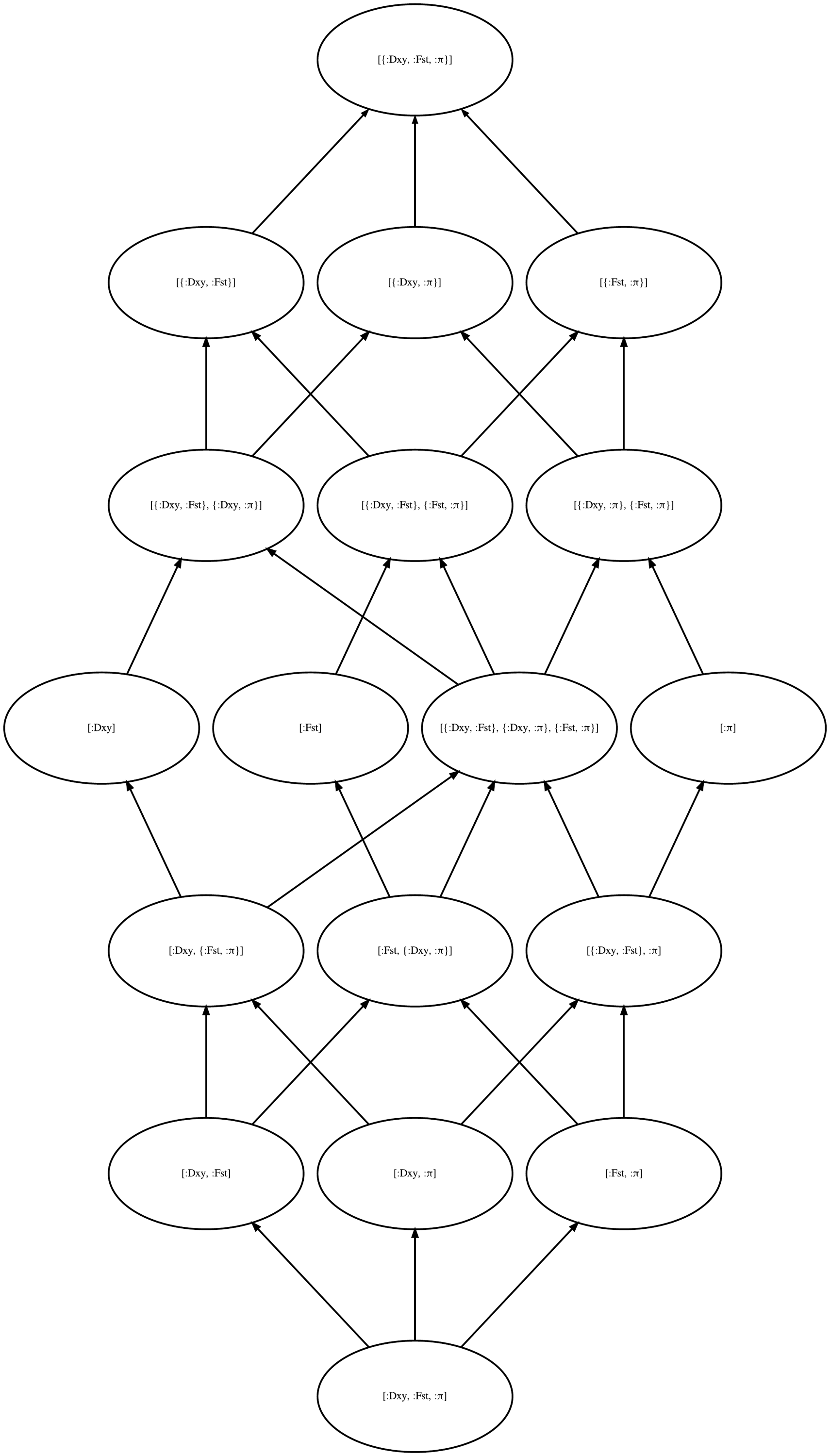

**Figure S11 Full decomposition lattice for [**D_XY_, F_ST_, π] **shown in Figure 3B.**

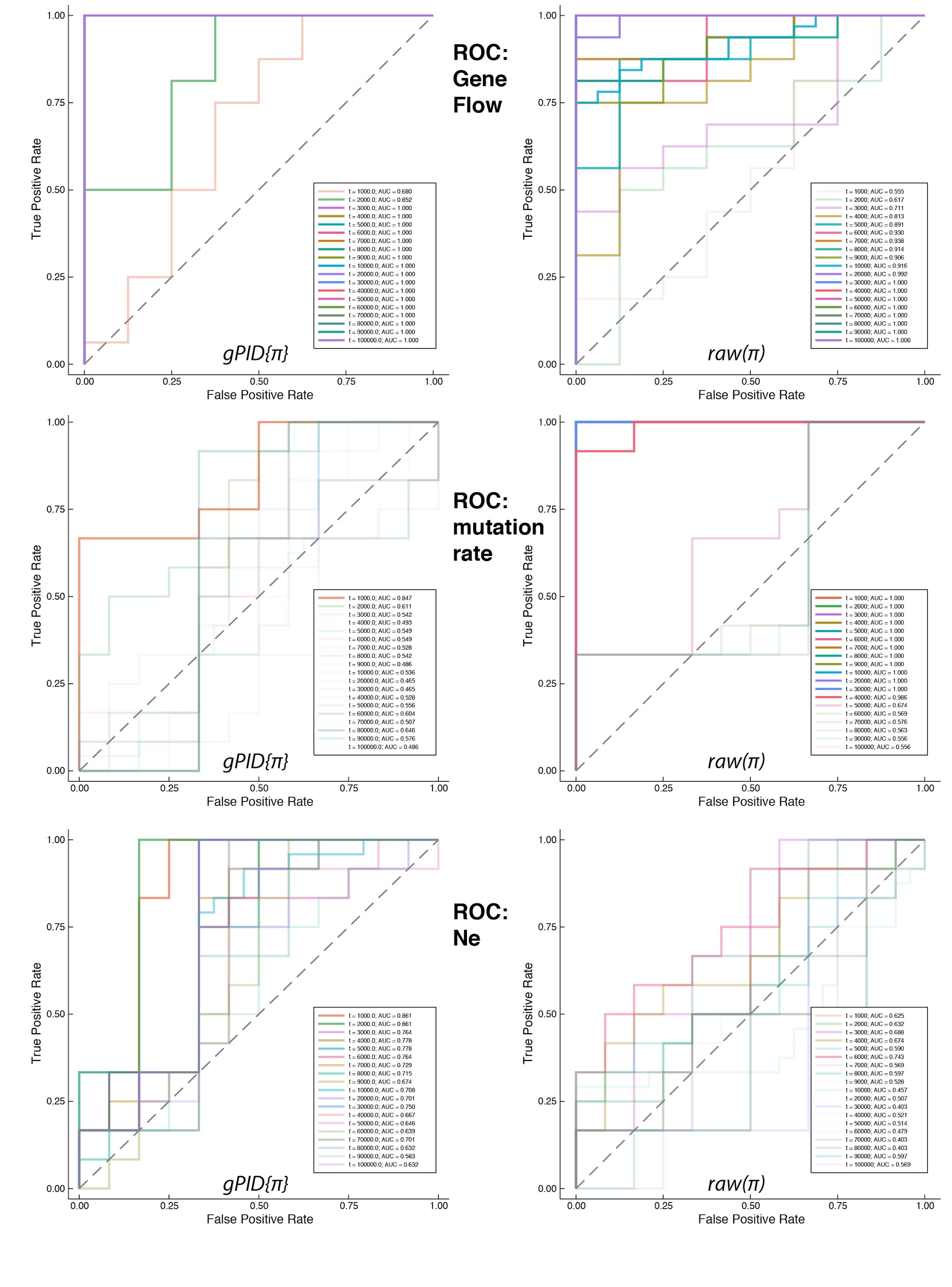

**Figure S12. Receiver-operator curves (ROCs) for the decomposition of partial information of 𝜋 (left) and associated raw values (right)** for three evolutionary parameters (top to bottom). The ROCs represent the ability of each variable to discriminate between datasets simulated under different values of that parameter (e.g., gene flow). Gene flow is the most widely detected signal (top), with some discrimination potential after more generations since divergence for both mutation rate and effective population size. PID performs better than using the raw values under all conditions except that within the first 1000 generations, raw values for 𝜋 can discriminate among mutation rates, but this is a temporary signal.

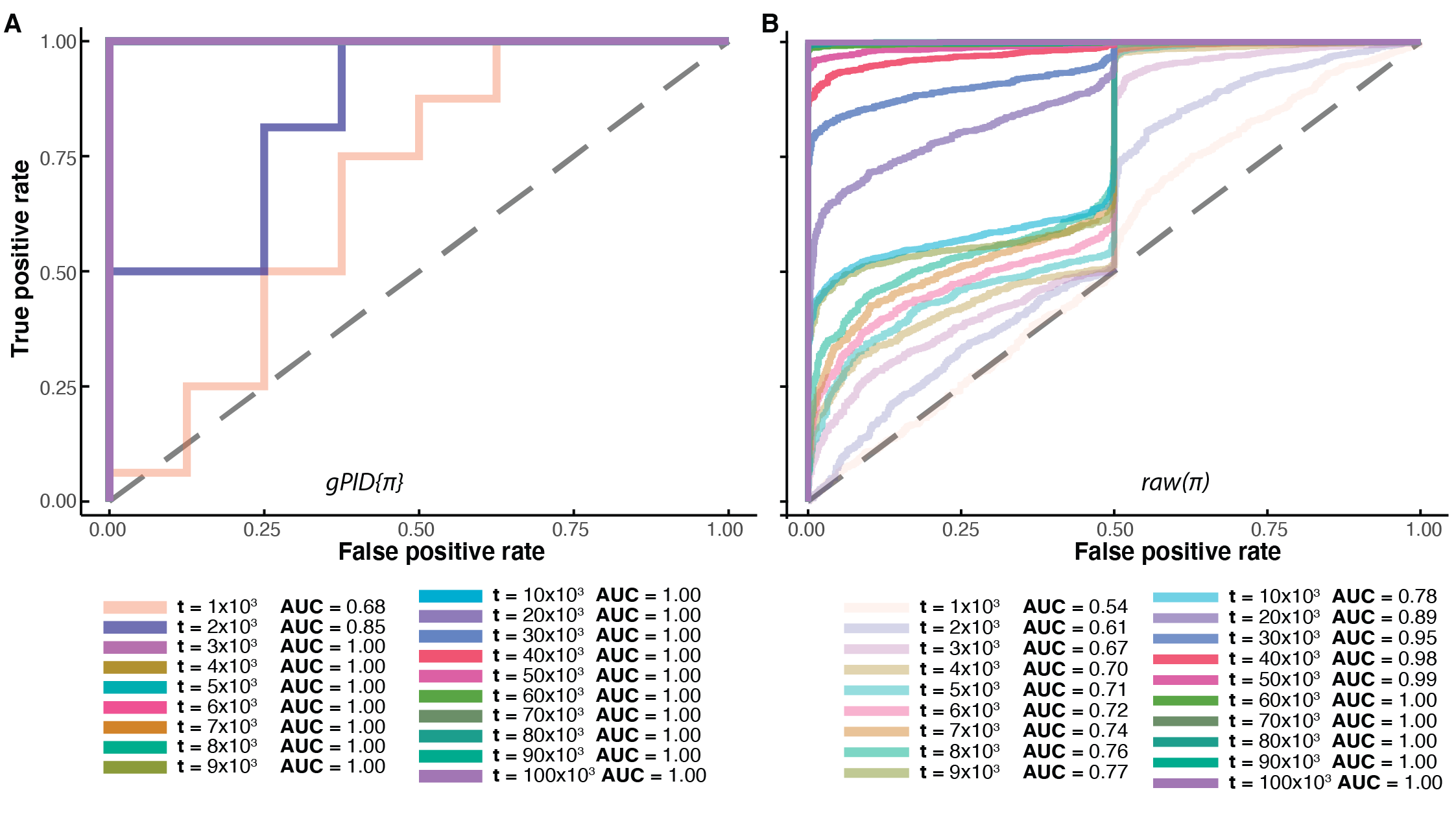

**Figure S13	Receiver-operating curve of nucleotide diversity (𝜋)** from a decomposed node (A) versus raw values of 𝜋 reflecting the ability for these metrics to discriminate between presence and absence of gene flow.
